# Plasticity and environmental heterogeneity predict geographic resilience patterns of foundation species to future change

**DOI:** 10.1101/401588

**Authors:** Luca Telesca, Lloyd S. Peck, Trystan Sanders, Jakob Thyrring, Mikael K. Sejr, Elizabeth M. Harper

**Affiliations:** Department of Earth Sciences, University of Cambridge, CB2 3EQ Cambridge, UK.; British Antarctic Survey, CB3 0ET Cambridge, UK.; GEOMAR Helmholtz Centre for Ocean Research, 24105 Kiel, Germany.; Department of Bioscience, Arctic Research Centre, Aarhus University, 8000 Aarhus C, Denmark.; Department of Bioscience, Marine Ecology, Aarhus University, 8600 Silkeborg, Denmark.

## Abstract

Although geographic patterns of species’ sensitivity to global environmental changes are defined by interacting multiple stressors, little is known about the biological mechanisms shaping regional differences in organismal vulnerability. Here, we examine large-scale spatial variations in biomineralisation under heterogeneous environmental gradients of temperature, salinity and food availability across a 30° latitudinal range (3,334 km), to test whether plasticity in calcareous shell production and composition, from juveniles to large adults, mediates geographic patterns of resilience to climate change in critical foundation species, the mussels *Mytilus edulis* and *M. trossulus*. We find mussels produced thinner shells with a higher organic content in polar than temperature regions, indicating decreasing shell calcification towards high latitudes. Salinity was the major driver of regional differences in mussel shell deposition, and in shell mineral and organic composition. In low-salinity environments, the production of calcite and organic shell layers was increased, providing higher resistance against dissolution in more corrosive waters. Conversely, under higher-salinity regimes, increased aragonite deposition suggests enhanced mechanical protection from predators. Interacting strong effects of decreasing salinity and increasing food availability on the compositional shell plasticity in polar and subpolar mussels during growth predict the deposition of a thicker external organic layer (periostracum) under forecasted future environmental conditions. This marked response potential of *Mytilus* species suggests a capacity for increased protection of high-latitude mussel populations from ocean acidification. Our work illustrates that mechanisms driving plastic responses to the spatial structure of multiple stressors can define geographic patterns of unforeseen species resilience to global environmental change.

## INTRODUCTION

Unprecedented global environmental changes are driving scientists towards increased understanding of the mechanisms underlying geographic variation in species’ responses to future environmental conditions (*1*, *2*). However, our ability to forecast emergent ecological consequences of climate change on marine populations, communities and ecosystems remains limited (*3*). Ecosystem-wide projections are severely constrained by heterogeneous patterns of ocean warming and acidification (*4*), multiple interacting stressors (*5*), and species-specific effects (*6*), as well as predictive models which often exclude important biological mechanisms when projecting changes to species and ecosystems in response to climate change (*2*). A better mechanistic understanding of the biological processes and environmental sources mediating species’ responses to disturbances is critical for building the theoretical baseline necessary to forecast the combined effects of multiple emerging stressors (*2*, *3*).

Advances in macroecology suggest that permanent environmental mosaics, defined by spatial overlaps of non-monotonic environmental gradients (*7*), as well as regional adaption or acclimatization (*8*–*10*), dictate geographic variations in species performance and sensitivity to environmental change in marine ecosystems. Key to these works is that responses vary among populations and individual taxa (*6*, *8*), which often play disproportionately strong roles in structuring benthic communities (*11*). Thus, species-specific biological mechanisms driving organismal variability may shape differential regional responses of foundation species to co-occurring multiple drivers. This can establish spatial patterns of unexpected susceptibility of marine communities to future conditions.

Climate change is considered a major threat to marine ecosystems worldwide, with ocean warming and acidification profoundly affecting species life history and ecology (*6*, *10*), as well as community structure and ecosystem dynamics (*11*, *12*). Species producing calcium carbonate (CaCO_3_) shells and skeletons are possibly experiencing the strongest impacts of rapid environmental changes (*6*). Knowledge on their sensitivity is derived largely from experimentally induced responses in model organisms (*1*, *6*), while complex variations under multiple stressors have rarely been investigated in natural environments (*7*, *11*–*13*). Therefore, inferences made from experimental studies can be misleading and not fully applicable to marine ecosystems (*9*). Indeed, species-specific mechanistic responses to habitat alterations (*14*) on top of mixed outcomes of environmental interactions (additive, synergistic or antagonistic) make future ecosystem predictions extremely challenging. This leaves open the question: do differences in biological mechanisms, shaping regional calcifiers’ responses to interacting environmental stressors, define geographic patterns of unforeseen species sensitivity or resilience to global environmental change? A body of research has focused on responses of marine calcifiers to altered water chemistry (*1*, *6*), but studies have rarely considered changes in biogeochemical cycles that strongly mediate biological responses to environmental alterations (*4*). Among those, a marked intensification of the global water cycle in response to warming (+4% for +0.5°C) has been documented over recent decades through changes in ocean salinity (*15*). Salinity is a major ecological factor dictating survival of aquatic organisms and ecosystem functioning. Multidecadal studies have revealed a global salinity pattern following the “rich-get-richer” mechanism, where salty ocean regions (compared to the global mean) are getting saltier (mid-latitudes), whereas low salinity regions are getting fresher (tropical convergence zones and polar regions) (*15*). In a future 2-3°C warmer world (*16*), a substantial 16-24% intensification of the global water cycle is predicted to occur making salinity gradients much sharper (*15*). However, emergent ecological effects of changing salinity on calcifying species and marine communities are largely unknown.

Atlantic mussels, *Mytilus edulis* and *M. trossulus*, are important bed-forming foundation species throughout the eulittoral ecosystems of the northern hemisphere (up to 90% of epibenthic biomass), and represent valuable resources for aquaculture (192,000 t produced in 2015 worth 325 million USD) (*17*). Growing awareness of the consequences of climate change on biodiversity and industry that *Mytilus* species support have stimulated a number of studies to estimate the response potential of these habitat-forming calcifiers to changing ocean conditions (*18*–*20*).

Calcareous shells perform a range of vital functions including structural support and protection against predators. Because shell integrity determines survival, shell traits are subject to strong selection pressure with functional success or failure a fundamental evolutionary driver. *Mytilus* shell consists of three layers (Fig. 1A-B): (1) the outer organic periostracum, (2) the calcified prismatic and (3) nacreous layers. The periostracum provides the substrate and a protected environment for shell secretion, and is made of sclerotized proteins, protecting shells from corrosive, acidic waters as well as predatory and endolithic borers (*21*). The prismatic and nacreous layers are composed of different mineral forms of CaCO_3_, calcite and aragonite respectively, dispersed in an organic matrix (*22*). These calcareous layers are characterized by different microstructures and more (e.g. aragonite) or less (e.g. calcite and organics) soluble components the combination of which determines chemical and mechanical shell properties (*23*). Differences in energetic costs of making shell components (*13*) combined with future shifts in environmental gradients (*4*) may influence variations in shell production, composition and structure, shaping regional patterns of shell strength and resistance to acidification.

**Fig. 1.**
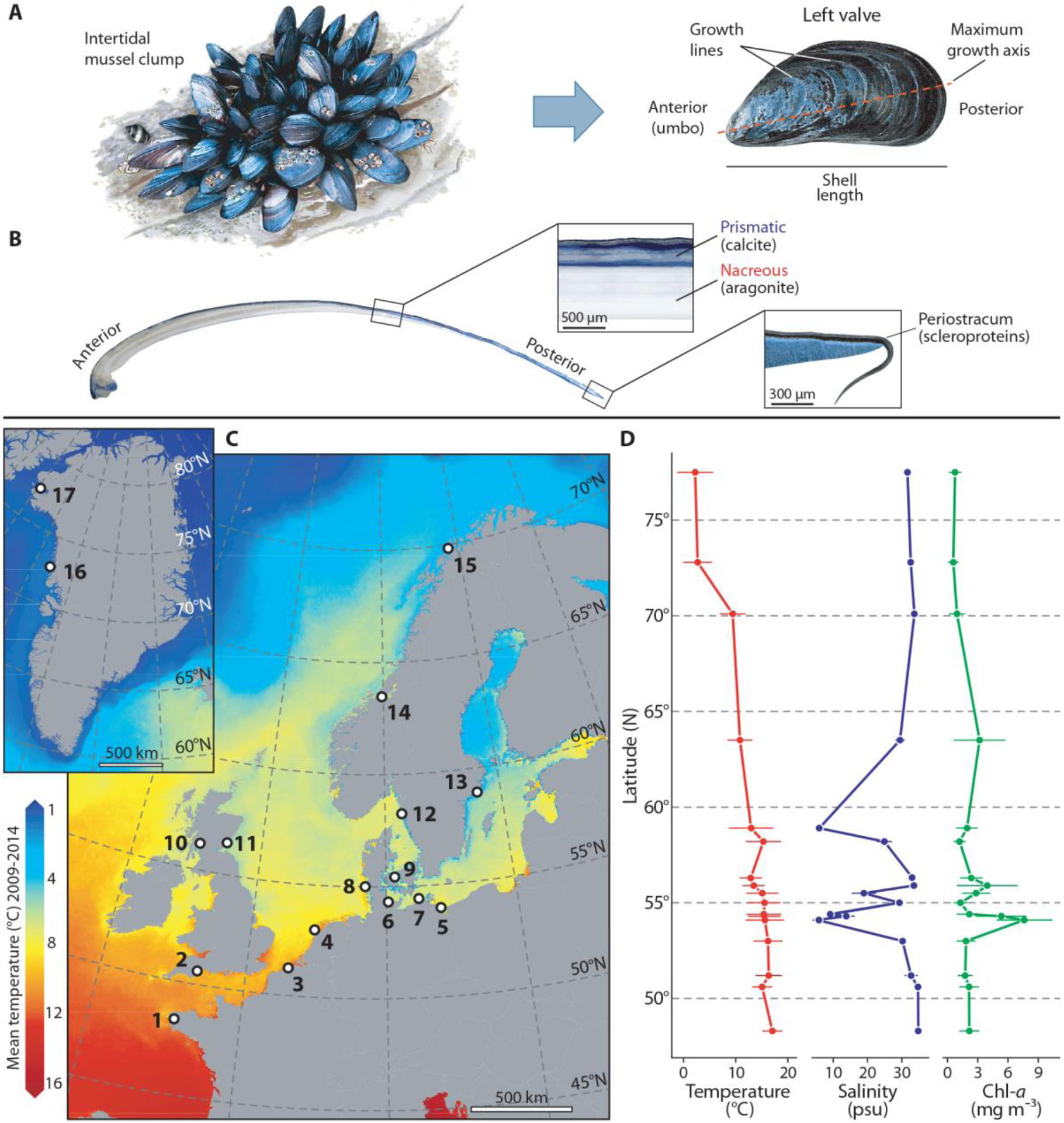
*Mytilus* spp. shell, collection sites and environmental heterogeneity. (**A**) *Mytilus* shell valve morphology and dimensions. (**B**) Anteroposterior cross-section of shell valve along the axis of maximum growth (from umbo to posterior commissure) showing internal structure and composition of individual mineral (prismatic and nacreous) and organic (periostracum) shell layers. (**C**) Thermal map of North-East Atlantic and Arctic surface waters from the CMEMS (http://marine.copernicus.eu/) biogeochemical datasets showing locations (open circles) where *Mytilus* was collected from across the Eastern European and Greenlandic coastlines (from 48°N to 78°N): (1) Brest, France, (2) Exmouth, England, (3) Oostende, Belgium, (4) Texel, Netherlands, (5) Usedom, (6) Kiel, (7) Ahrenshoop, (8) Sylt, all Germany, (9) Kerteminde, Denmark, (10) Tarbet, Scotland, (11) St. Andrews, Scotland, (12) Kristineberg, Sweden, (13) Nynäshamn, Sweden (14) Trondhiem, Norway, (15) Tromsø, Norway, (16) Upernavik, Greenland and (17) Qaanaaq, Greenland. Map created with ArcMap 10.5 (ArcGIS software by Esri, http://esri.com), background image courtesy of OpenStreetMap (http://www.openstreetmap.org). (**D**) Latitudinal gradients for sea surface temperature, salinity and chlorophyll-*a* (Chl-*a*) concentration, showing environmental heterogeneity across the study regions. Mean values (May - October, filled circles) and SD (horizontal lines) for the 6-year period 2009 - 2014 were estimated from CMEMS datasets.

*Mytilus* growth, biomineralisation and fitness are linked to multiple drivers, including water temperature, salinity and food supply [chlorophyll-*a* (Chl-*a*) concentration] (*24*, *25*). In the North Atlantic and Arctic Oceans, these key environmental factors vary heterogeneously with latitude (Fig. 1C-D), encompassing a range of conditions predicted under different future climate change scenarios (*16*). Here we hypothesize that biological mechanisms driving spatial variations in shell production, mineral and organic composition: **i**) shape regional differences in the responses of *Mytilus* species to interacting environmental drivers, and **ii**) define geographic patterns of unanticipated mussel vulnerability in the face of global environmental changes.

Despite projected environmental alterations (*4*, *15*), salinity gradients have been overlooked in large-scale models predicting emergent effects of climate changes on marine organisms. This knowledge is essential to predict whether environmental changes affect shell variability (i.e. thickness, mineral and organic content) and its properties, especially in calcifying foundation species such as *M. edulis* and *M. trossulus*. These factors are crucial for understanding species susceptibility to other rapidly emergent stressors, such as warming and acidification (*3*).

In this study, we examine the relationships between the plasticity in *Mytilus* shell production and composition (from juveniles to large adults) and interactive environmental gradients of temperature, salinity and Chl-*a* concentration in 17 populations spanning a latitudinal range of 30° (3,334 km) across the Atlantic-European and Arctic coastline (Fig. 1C-D). In particular, we test for a latitudinal effect on *Mytilus* shell calcification (variation in shell thickness and organic content) that we hypothesize will show a general decrease from temperate to polar regions. We also identified environmental sources and magnitude of regional variations in shell deposition, to test whether salinity affects shell production and mineral composition during growth, driving changes of mechanical and chemical shell properties. Finally, we modelled spatial trends in the production of individual shell layers with environmental gradients, to test whether biological mechanisms, driving variations in shell structure and properties, shape regional responses of *Mytilus* to interacting stressors (especially salinity) and define geographic patterns of sensitivity to future changes.

## RESULTS

Generalized linear (mixed) models, GL(M)Ms, were used to explain shell thickness and composition, from juveniles to large adults, with respect to latitude and environmental drivers, and to compare between the individual shell layers.

### Latitudinal patterns of shell deposition

GLMMs indicated a general decrease of *Mytilus* whole-shell thickness with increasing latitude from warm-temperate to polar regions (Fig. 2A). We detected a significant negative relationship between the prismatic layer thickness and latitude (Fig. 2A), while no variation in nacreous thickness, periostracum thickness and relative proportion of prismatic layer thickness (calcite%) was found (table S1). Shell length was positively correlated with thickness in all layers indicating thickening during growth (table S1).

**Fig. 2.**
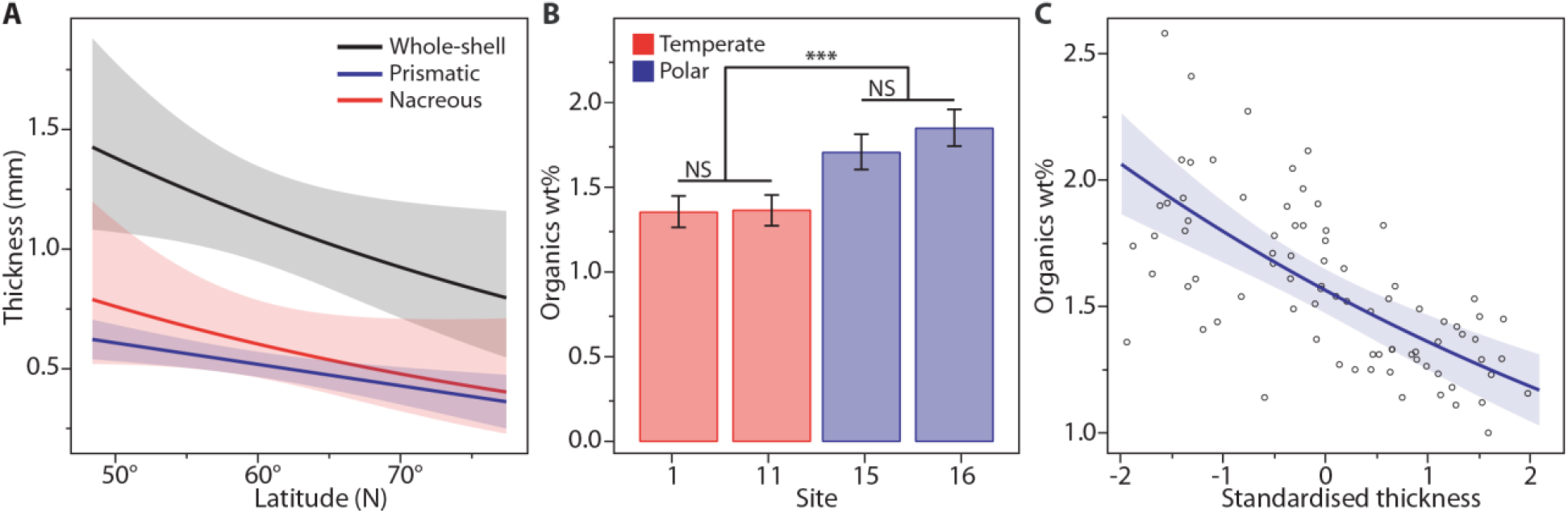
Latitudinal patterns of shell thickness, organic content and calcification. (**A**) Relationships between the thickness of whole-shell (black), prismatic (blue) and nacreous (red) layers and latitude. Whole-shell thickness decreased poleward (95% CI = -0.36 to - 0.01, cR^2^ = 0.81). The prismatic layer was significantly related to latitude (95% CI = 4.70 to 5.73, cR^2^ = 0.72). Predicted values (continuous lines) and confidence intervals (shaded areas) were estimated for mussels of mean shell length (47.42 mm). Parameters’ significance is determined when the bootstrapped 95% CI does not include zero. (**B**) Variations in organic content among shells from temperate (sites 1, 11, white bars) and polar (sites 15, 16, grey bars) climates. Pair-wise contrasts indicated significantly higher proportions of organics in high-latitude than low-latitude specimens [mean difference = 0.44%; *z* = 8.27, *P* < 0.0001 (***), pseudoR^2^ = 0.49], in addition to non-significant differences (NS) among temperate (mean difference = 0.002%; *z* = 0.12, *P* = 0.91) and polar (mean difference = 0.13%, *z* = 1.86, *P* = 0.063) populations. (**C**) Relationship between the proportion of organics and standardised thickness of the prismatic [mean (SD) = 529 µm (174)] (sites 1, 7, 10 and 11), indicating a negative association between layer thickness and calcification level (*z* = -7.10, *P* < 0.0001, pseudoR^2^ = 0.40).

The weight proportion (wt%) of organic content in the prismatic layer was modelled with a GLM as a function of collection site and shell thickness. Prismatic layers were characterized by a significantly higher organic content (lower proportion of CaCO_3_) in mussel shells from polar than temperate regions, indicating decreased shell calcification at higher latitudes (Fig. 2B). Polar shells [sites 15, 16; mean (SD) = 1.8 wt% (0.31)] were characterized by an average of 29% more organic content compared to temperate mussels [sites 1, 11; mean (SD) = 1.4 wt% (0.16)]. The organics wt% was negatively correlated with prismatic thickness (Fig. 2C), indicating a lower proportion of CaCO_3_ and thinner, so less calcified, shells at polar latitudes.

### Environmental influence on shell production and composition

Individual GLMMs were fitted to explain spatial variations in the whole-shell thickness, periostracum thickness and calcite% with environmental gradients during shell growth. We identified significant trends in shell thickness with environmental gradients depending on the shell measurement considered (Fig. 3A-E, fig. S1, Table 1). Whole-shell thickness was positively related to temperature, salinity and shell length, but there was no influence of Chl-*a* (cR^2^ = 0.93; Fig. 3A). Salinity had an effect on shell thickness that was 3.4 and 2.1 times larger than temperature and length, respectively (Fig. 3A, Table 1). We detected a negative relationship between calcite% and salinity (95% CI = -12.03 to -2.38, cR^2^ = 0.56) (Fig. 3B, table S2), with none of the other drivers having a significant effect.

**Fig. 3.**
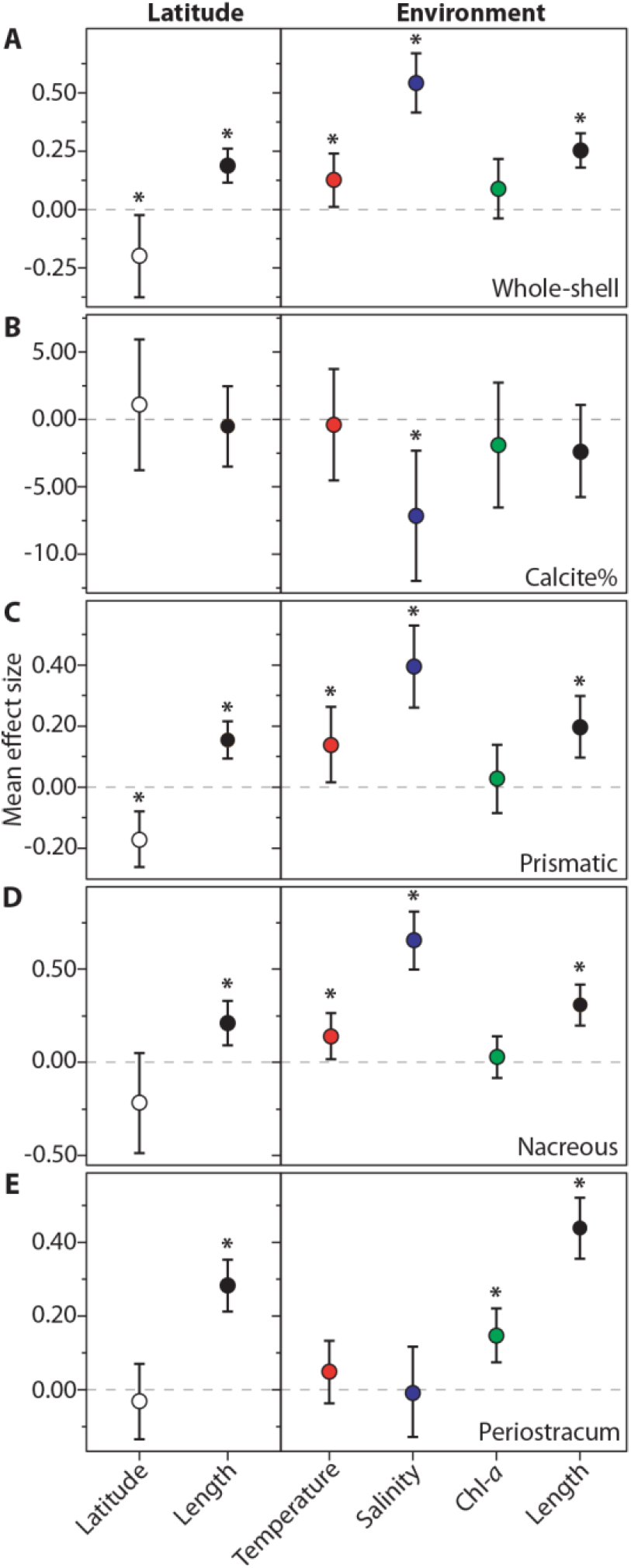
Mean effect size of predictors on *Mytilus* shell measurements. Effect sizes were estimated from individual latitudinal (left panels) and environmental (right panels) GLMMs. Mean effect sizes and direction of impacts of latitude (white), shell length (black), sea surface temperature (red), salinity (blue) and Chl-*a* concentration (green) on layer ln-thickness (µm) measurements and calcite% are reported: (**A**) whole-shell, (**B**) calcite%, (**C**) prismatic layer, (**D**) nacreous layer and (**E**) periostracum. Note the different scales on the y-axis to highlight variations among layers. Significance of regression parameters is determined when the bootstrapped 95% CI (error bars) does not cross zero (* denotes a significant difference from zero).

**Table 1.** Environmental GLMMs summary. Estimated statistics and bootstrapped 95% CIs for regression parameters are reported for the modelled relationships between individual shell thickness measurements and standardised covariates. For the summary of model in equation (1), the prismatic layer, Layer(Pr), is used as the reference level, (Intercept). (Parameters’ significance is determined when the 95% CI does not include zero).

|  | Estimate | SE | 95% CI | *t*-value | *P*-value<br>(approximate) |
| --- | --- | --- | --- | --- | --- |
| **Whole-shell\*** |  |  |  |  |  |
| (Intercept) | 6.617 | 0.051 | 6.517; 6.717 | 128.71 | <0.0001 |
| Temperature | 0.156 | 0.054 | 0.014; 0.240 | 2.89 | 0.013 |
| Salinity | 0.525 | 0.060 | 0.411; 0.672 | 8.69 | <0.0001 |
| Chl-*a* | 0.074 | 0.054 | -0.042; 0.216 | 1.37 | 0.20 |
| Length | 0.248 | 0.037 | 0.181; 0.327 | 6.44 | <0.0001 |
| **Prismatic (Pr) & nacreous (Na)<sup>†</sup>** |  |  |  |  |  |
| (Intercept) | 5.907 | 0.031 | 5.774; 6.036 | 188.31 | <0.0001 |
| Temperature | 0.138 | 0.033 | 0.013; 0.260 | 4.17 | 0.0011 |
| Salinity | 0.396 | 0.039 | 0.264; 0.531 | 10.22 | <0.0001 |
| Chl-*a* | 0.028 | 0.033 | -0.087; 0.139 | 0.86 | 0.41 |
| Length | 0.197 | 0.031 | 0.096; 0.297 | 6.39 | <0.0001 |
| Layer(Na) | -0.054 | 0.078 | -0.247; 0.134 | -0.70 | 0.49 |
| Salinity ×<br>Layer(Na) | 0.259 | 0.088 | 0.065; 0.443 | 2.95 | 0.0033 |
| Length ×<br>Layer(Na) | 0.111 | 0.067 | -0.036; 0.258 | 1.66 | 0.096 |
| **Periostracum<sup>‡</sup>** |  |  |  |  |  |
| (Intercept) | 3.500 | 0.048 | 3.406; 3.596 | 71.03 | <0.0001 |
| Temperature | 0.049 | 0.043 | -0.036; 0.134 | 1.12 | 0.28 |
| Salinity | -0.009 | 0.061 | -0.131; 0.111 | -0.14 | 0.89 |
| Chl-*a* | 0.147 | 0.038 | 0.071; 0.221 | 3.88 | 0.0020 |
| Length | 0.439 | 0.041 | 0.357; 0.522 | 10.25 | <0.0001 |
| Temperature ×<br>Length | -0.064 | 0.035 | -0.135; 0.006 | -1.77 | 0.082 |
| Salinity ×<br>Length | -0.151 | 0.061 | -0.271; -0.029 | -2.38 | 0.020 |

\* Whole-shell, the random intercept was normally distributed with mean of 0 and variance 0.209<sup>2</sup>.
<sup>†</sup> Prismatic and nacreous layer, the random intercepts were normally distributed with mean 0, and variances 0.123<sup>2</sup> and 0.310<sup>2</sup>, respectively.
<sup>‡</sup> Periostracum, the random intercept was normally distributed with mean 0 and variance 0.130<sup>2</sup>.

Prismatic and nacreous layers thickness were analysed within the same GLMM. After model selection, fixed continuous covariates of the optimal model, equation (1), were the standardised *temperature*, *salinity*, *Chl-a* concentration and shell *length* in addition to shell *layer* (categorical, two levels: prismatic and nacreous) and their interactions. The random component was the collection *site* used as a random intercept. The model was of the form:

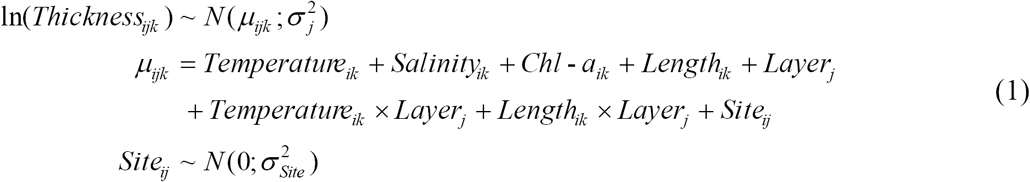

where *Thickness*_*ijk*_ is the *k*th thickness observation from layer *j* (*j* = prismatic, nacreous) and site *i* (*i* = 1,…, 17). *Site*_*ij*_ is the random intercept for layer *j*, which is assumed to be normally distributed with mean 0 and variance 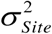

Sea surface temperature, salinity and shell length all successfully predicted (cR^2^ = 0.93) variations in the thickness of prismatic and nacreous layers, while no influence of Chl-*a* was detected (Table 1). The mean effect size of salinity on the response was twice as large as the effect of shell length, while it was 2.9 and 4.7 times larger than the effect of temperature on the prismatic and nacreous layers, respectively (equation (2), Fig. 3C-D). This indicates salinity had a stronger contribution to predicting shell structure than the effects of temperature, Chl-*a* and shell length combined (Fig. 4).

**Fig. 4.**
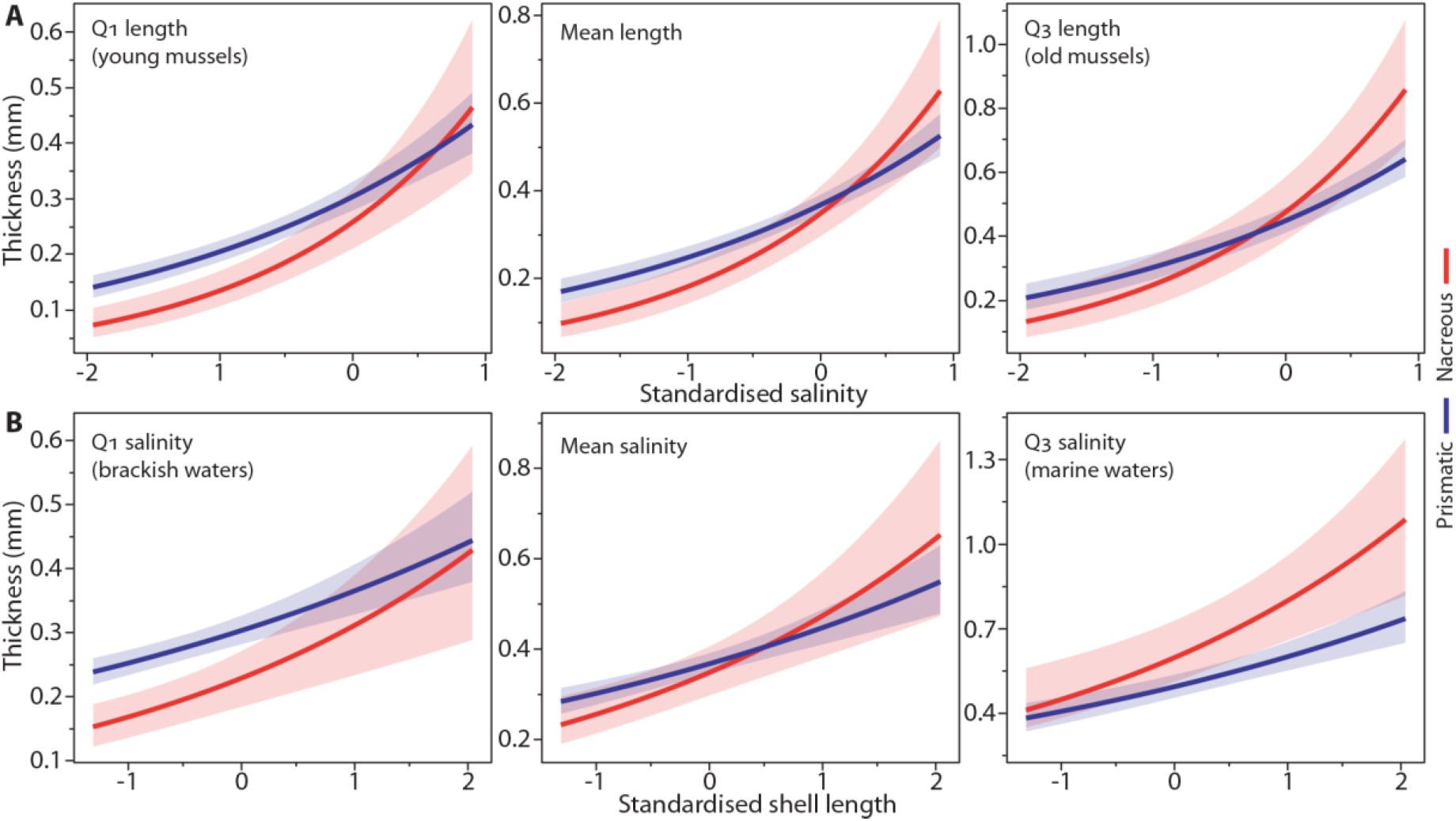
Environmental influence on shell production and composition. Predicted relationships between thickness of prismatic (blue) and nacreous (red) layers, and standardised salinity [mean (SD) = 25.52 psu (10.29)], shell length [mean (SD) = 47.42 mm (16.20)] and their interactions. (**A**) Shell thickness is modelled as a function of salinity for the 1st quartile (Q_1_ = 31.50 mm), mean value (47.42 mm) and 3rd quartile (Q_3_ = 63.90 mm) of the shell lengths sampled. For medium-sized mussels, we detected a decreasing proportion of the calcitic component with increasing salinity and the deposition of relatively thicker aragonitic layers at salinities > 27.67 psu. (**B**) Thickness is modelled as a function of length for the 1st quartile (Q_1_ = 18.92 psu), mean value (25.52 psu) and 3rd quartile (Q_3_ = 33.13 psu) of salinity. At mean salinity, we detected an inversion of the relative layers’ thickness for shell length > 55.30 mm. Across the entire range of shell lengths, the model predicts formation of calcite- and nacreous-dominated shells under low- and high-salinities, respectively. Mean values (continuous lines) and confidence intervals (shaded areas) are predicted controlling for temperature (13.03°C) and Chl-*a* (2.48 mg m^-3^).

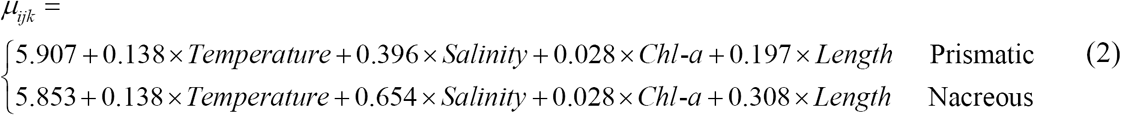

Interactions between shell layer and both salinity and shell length (equation (2)) indicate deposition of proportionally thicker prismatic layers under low salinities and proportionally thicker nacreous layers under higher salinities across the entire range of shell lengths (Fig. 4).

### Periostracum plasticity

Models of periostracum thickness revealed significant exponential relationships with Chl-*a* and shell length (cR^2^ = 0.81) (Table 1). Length had a mean effect that was 3 times larger than Chl-*a* (Fig. 3E), showing a rapid thickening of the periostracum during shell growth. The interactions between shell length and both salinity and temperature indicate that the effects of these variables on periostracum were interdependent. At low salinities, the higher values of shell length had a greater positive effect on periostracum thickness, while the reverse was true for higher temperatures having a marginal effect only on thickening rates (Fig. 5A-B). This suggests that increasing shell size was a more important factor for periostracum growth in fresher waters than in relatively saltier conditions.

**Fig. 5.**
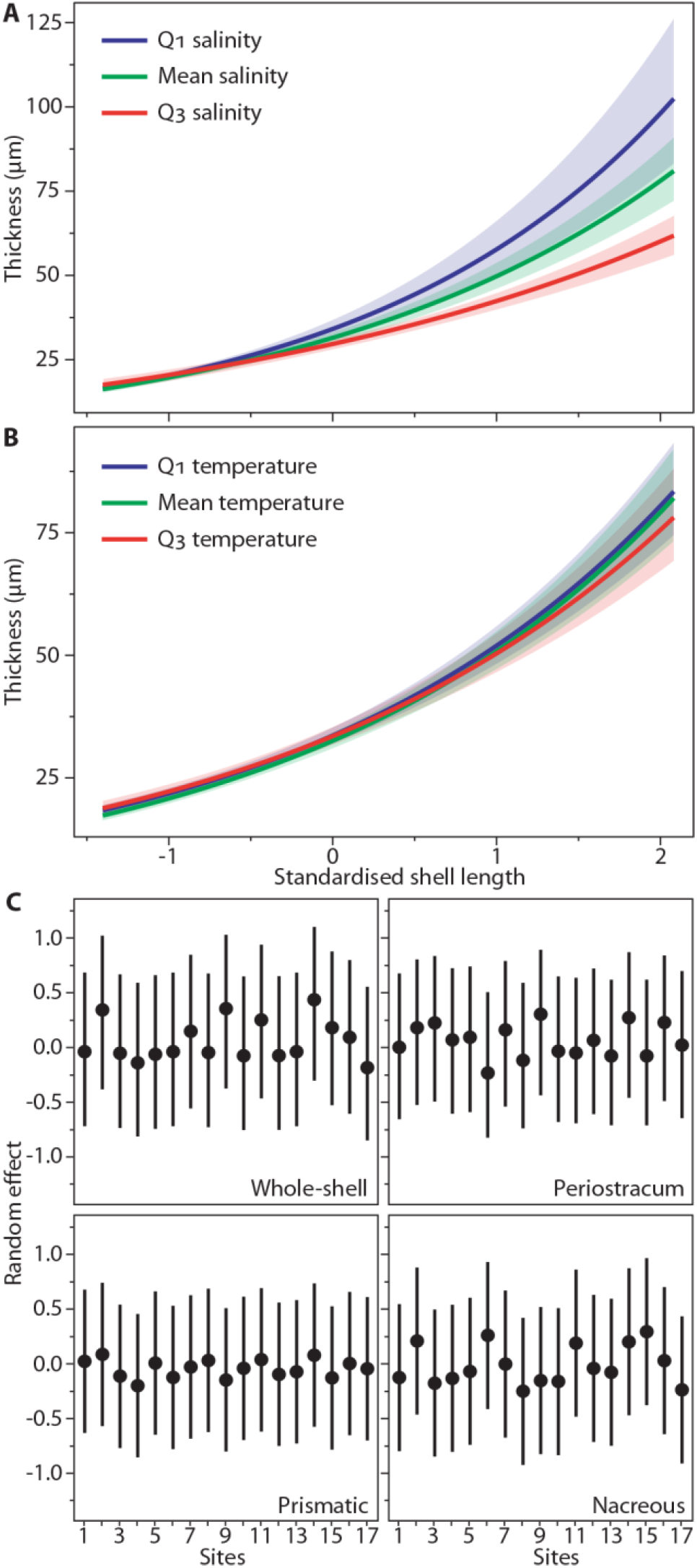
Periostracum plasticity and among-site shell variation. Interacting effects of salinity, temperature and shell length on shell periostracum. (**A**) Periostracum thickness is modelled as a function of shell length [mean (SD) = 47.42 mm (16.20)] for the 1st quartile (Q_1_ = 18.92 psu, blue line), mean (25.52 psu, black line) and 3rd quartile (Q_3_ = 33.13 psu, red line) of water salinity. Predicted values (continuous lines) and confidence intervals (shaded areas) indicate higher rates of exponential periostracal thickening with decreasing salinity. Smaller individuals (shell length < 48.38 mm) were characterized by non-significant thickness differences under different salinity regimes. (**B**) Thickness is modelled for the 1st quartile (Q_1_ = 12.89°C, blue line), mean (13.03°C, black line) and 3rd quartile (Q_3_ = 15.51°C, red line) of water temperature. Predicted values indicate a marginal influence of temperature on periostracal thickening. (**C**) GLMMs’ conditional modes (filled circles) and variances (continuous lines) of the random effect estimated for individual shell layers. Modes represent the difference between the average predicted response (layer thickness) for a given set of fixed-effects values (mean environmental covariates and shell length) and the response predicted at a particular site. These indicate no detectable residual effect of species (*Mytilus edulis* or *M. trossulus*) and level of hybridization on shell thickness for each site.

### Among-site shell variation

GLMMs showed no difference in collection site-level effects (conditional modes) on each thickness measurement (Fig. 5C). This indicated no residual effect of species identity or hybridization on the thickness of individual shell layers at different sites after accounting for the effects of environmental factors and shell length.

## DISCUSSION

Our results demonstrate that plasticity in shell production in *Mytilus* species and the spatial structure of environmental conditions drive geographic variations in shell responses shaping regional differences in the resilience of these foundation species to global environmental change. An understanding of the biological mechanisms driving regional species’ responses to multiple interacting stressors is crucial for improving predictive accuracy and informing more realistic projections of species and ecosystem resilience to climate change (*2*). Heterogeneous population-level responses from different climates suggest that environmental stressors, especially salinity, drive regional variations in *Mytilus* shell production, mineral (prismatic and nacreous) and organic (periostracum) composition during growth, which is reflected in the relative proportion of each shell layer. Variations in shell production and composition determine geographic differences in chemical and mechanical protection of shells, shaping the vulnerability of these habitat-forming species to future conditions.

Decreasing shell calcification (increasing organic content and thinner shells) towards high latitudes (Fig. 2) supports documented patterns of skeletal production (*13*, *26*). Two explanatory paradigms exist for decreased skeletal size at higher latitudes: increased calcification costs (*13*) and reduced predation pressure (*27*). Given the higher production cost of organics than CaCO_3_ deposition (*13*) and problematic protein production at polar temperatures (*28*), we might expect a reduced proportion of organic matrix. Moreover, decreasing predation pressure (*27*) should result in thinner shells of the same composition irrespective of geographic area. However, the wt% of organic matrix was higher at Arctic latitudes. This could suggest either (or a combination of) a marked increase in the cost of calcification in polar regions (*13*), altering significantly the relative costs of CaCO_3_ and organics production, or a decreased saturation state (increased dissolution) of CaCO_3_ due to low temperatures and, more importantly, salinity (low [Ca^2+^] availability) (*25*). In either case, these underlying effects would result in decreased shell calcification at high latitudes. This increased proportion of organic matrix could protect the calcified shell components from dissolution and have an adaptive beneficial effect in more corrosive conditions.

Our results illustrate that different drivers significantly affect both shell thickness and composition in *Mytilus* (Fig. 3). For over 60 years, temperature and shell size have been considered key drivers of CaCO_3_ shell mineralogy across latitudes, dictating the formation of predominantly aragonitic structures in temperate regions and increased calcite precipitation in cold climates (*29*–*31*). Although our study partly supports previous findings, we demonstrate that salinity has the strongest influence on shell production and composition in *Mytilus*, which is contrary to the general assumption of temperature and shell size being the primary drivers of shell compositional plasticity.

The interaction between shell layer, salinity and shell size indicates heterogeneous, age-related compositional changes in *Mytilus* shells across different salinities (Fig. 4A). Shifts in shell properties from juveniles to large adults are strongly modulated by salinity, which leads to the formation of exclusively prismatic-dominated shells in brackish waters and nacreous-dominated structures under marine conditions (Fig. 4B). These patterns, which we show were independent of species or hybrid status (Fig. 5C), indicate that mussel shell plasticity during growth (the *Length* × *Layer* interaction, equation (1)) has an indirect effect on layer thickness by allowing salinity-induced compositional changes and, therefore, the production of the most appropriate shell structure for specific environmental conditions.

Under current scenarios, plasticity in shell production could confer *Mytilus* species an advantage when facing different water chemistries and predation levels. In fact, at high-latitudes and in the Baltic region, where durophagous (shell-breaking) predators are rare or absent and the water is more corrosive (*13*, *27*), mussels are characterized by thinner, prismatic-dominated shells, providing a generally higher protection from dissolution. Conversely, at mid-latitudes, where durophagous predators are more abundant and the CaCO_3_ solubility of the water is lower (*13*), mussels display thicker, nacreous-dominated shells with higher mechanical resistance.

Despite rapid global changes in the water cycle and salinity gradients (*15*), *Mytilus* species shows a strong capacity to respond to heterogeneous environments. This plasticity in shell production could help to mitigate the emergent negative effects of changing water chemistry. In fact, the interacting effects of salinity and shell length, as well as a minor influence of temperature, on the periostracum (Fig. 5A-B), which represents a strong chemical barrier to dissolution in molluscs (*21*, *32*, *33*), suggest enhanced periostracal thickness under decreasing salinities could mediate impacts of ocean acidification.

Although populations in high-latitude ecosystems will experience globally the most rapid acidification (*4*), the concurrent decrease in salinity predicts thicker prismatic layers and periostraca will be produced which increase protection from higher solubility conditions. Conversely, in temperate areas, increasing salinity would determine deposition of thicker shells and a relatively thicker nacreous layer and thinner periostracum, favouring mechanical shell resistance. However, predicted changes in periostracal thickening rate under different salinities depend on shell size and would be more evident in larger individuals (length > 48 mm) (Fig. 5A).

In Greenland, where the rate of melting of the ice sheet has doubled in the last decade (*34*), low salinities during summer (< 20 psu) and high productivity (food supply) in coastal areas and fjords (*35*) predict formation of thicker periostraca and increased relative thickness of organic-enriched (high wt%) calcitic layers. These changes could make Arctic *Mytilus* populations more resilient to future acidification. Differently, in the Baltic Sea, the forecasted decrease in salinity (maximum 45% reduction) (*36*), combined with a considerable physiological stress, would be particularly critical for mussels inhabiting already unfavourable conditions for calcification (salinity from 22 psu to 3 psu, low water [Ca^2+^], and CaCO_3_ saturation state) (*25*). Moreover, the reduced shell size of Baltic *Mytilus* does not predict formation of thicker periostraca, which would further increase vulnerability to dissolution. Impacts of changing salinity on this habitat-forming species, which contributes up to 90% of the Baltic benthic biomass, could strongly affect the ecosystem, most likely resulting in substantial range restrictions towards higher salinity areas.

Although our results strongly support the hypothesis that biological mechanisms for variations in shell production can shape regional responses in *Mytilus*, changes of other biological drivers, such as predation pressure and primary production, could also have profound influences (*13*, *37*). In fact, as temperature rises, durophagous predators could expand their ranges towards polar regions (*38*), suggesting an increased vulnerability of thin-shelled individuals. However, predicted northward phytoplankton expansions and an overall increase in primary production at high latitudes (*37*), could favour periostracal growth potential in *Mytilus* and, thus, increased resistance to dissolution for all the size classes in polar and subpolar regions.

*Mytilus* shells have a thick periostracum and a marked compositional plasticity compared to other calcifiers that often compete with it for space (e.g. barnacles and spirorbid polychaetes). This layer provides a strong defence against shell dissolution allowing mytilids to survive in oligohaline waters (∼5 psu) and extremely acidified conditions (e.g. hydrothermal vents) (*33*). These factors may shift the ecological balance and community structure in favour of species with stronger resistance to corrosive conditions, such as mussels, when ocean waters become fresher and more acidic in future decades.

As hypothesised, plasticity in shell production and the spatial structure of environmental conditions drive regional differences in *Mytilus* shell deposition and composition, shaping spatial patterns of chemical and mechanical shell properties. Overall, mussel shell calcification decreased towards high latitudes, with salinity being the major driver of geographical variations in shell production, mineral and organic composition. The marked compositional plasticity in calcareous shell components (prismatics and nacreous layers) suggest an higher resistance against dissolution for mussels in polar, low-salinity environments, and an enhanced mechanical shell protection from predators in temperate, higher-salinity regions. The strong response potential of *Mytilus* shell periostracum to heterogeneous environments predicts an increased resilience to ocean acidification in polar and sub-polar mussels, and a higher sensitivity of Baltic populations under future environmental conditions.

In conclusion, our findings demonstrate that biological mechanisms, driving spatial variability of mussel responses to interacting environmental factors, shape the complex geographic pattern of shell deposition and properties, dictating regional differences in *Mytilus* species sensitivity to future environmental change. As the magnitude of anthropogenic impacts continue to increase, further studies are need to better understand the key biological processes mediating species’ response to habitat alterations, especially for those having both high climate sensitivity and disproportionately strong ecological impacts in shaping marine communities. This knowledge underpins our ability to predict accurately and reduce the damaging effect of climate change on future biodiversity under any range of scenarios (*2*). Our study has important implications because it clarifies the links between (**i**) the mechanisms of biological variation, (**ii**) the predicted shift in spatial co-occurrence of multiple environmental drivers, and (**iii**) regional differences in the plastic responses and sensitivity of calcifying, foundation species to changing habitats. This understanding is of critical importance for making realistic projections of emergent ecological effects of global environmental changes, such as altered salinity regimes, and to improve our predictive accuracy for impacts on marine communities and ecosystems, and the services they provide.

## MATERIALS AND METHODS

### *Mytilus* collection

We sampled individuals from a total of 17 *Mytilus* (*Mytilus edulis* and *M. trossulus*) populations along the North Atlantic, Arctic and Baltic Sea coastlines from four distinctive climatic regions (warm-temperate, cold-temperate, subpolar and polar) covering a latitudinal range of 30° (a distance of 3,334 km), from Western European (Brest, North-West France, 48°N) to Northern Greenlandic (Qaanaaq, North-West Greenland 78°N) coastlines (Fig. 1C). During December 2014 - September 2015, mussels of various size classes for each site (shell length 26-81 mm) were sampled from the eulittoral zone on rocky shores for a total of 424 individuals (table S3). For each specimen, shell length was measured with digital calipers (0.01 mm precision) and used as a within-population proxy for age.

We analysed *Mytilus* populations of which the genetic structure was known, with particular focus on species identity and hybrid status (*M. edulis* × *M. trossulus*). *Mytilus* shells used were either from specimens already evaluated in genetic investigations or mussels obtained from sites routinely used in regional monitoring programs that provided information on species identity (table S3). Areas where the Mediterranean mussel, *Mytilus galloprovincialis*, was present were avoided. We did, however, sample few sites with very low levels of *M. edulis* × *M. galloprovincialis* hybridization.

### Mussel shell preparation

We set left shell valves in polyester resin (Kleer-Set FF, MetPrep, Coventry, U.K.) blocks. Embedded specimens were sliced longitudinally along their axis of maximum growth (Fig. 1A) using a diamond saw and then progressively polished with silicon carbide paper (grit size: P800-P2500) and diamond paste (grading: 9-1 µm). Photographs of polished sections (Fig. 1B) were acquired with a stereo-microscope (Leica M165 C equipped with a DFC295 HD camera, Leica, Wetzlar, Germany) and shell thickness (µm) was measured using the Fiji software (v1.51u). Since larger individuals had undergone evident environmental abrasion or dissolution which removed the periostracum and prismatic layer closer to the umbo, we estimated the thickness of the whole-shell, prismatic and nacreous layers at the midpoint along the shell cross-section. The proportion of calcite was estimated as

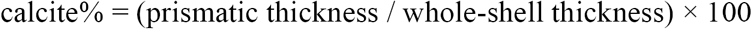

Periostracum thickness was measured at the posterior edge where it attaches to the external side of the prismatic layer, to estimate the fully formed organic layer that was unaffected by decay or abrasion (*21*).

### Organic content analyses

We performed thermogravimetric analyses (TGA) to estimate the weight proportion (wt%) of organic matrix within the prismatic layer. *Mytilus edulis* specimens were selected from four populations (sites 1, 11, 15, 16) to explore differences in shell organic content under temperate and polar regimes. We removed the periostracum by sanding, and prismatic layer tiles (8 × 5 mm, *N* = 20 × 4 sites) were isolated along the posteroventral shell margin. Tiles were cleaned, air-dried and then finely ground. We tested ten milligrams of this powdered shell with a thermogravimetric analyser (TGA Q500, TA Instruments, New Castle, DE, U.S.A.). Samples were subjected to constant heating from ∼25°C to 700°C at a linear rate of 10°C min^-1^ under a dynamic nitrogen atmosphere and weight changes were recorded (Supplementary Material and Methods). We estimated the wt% of organic matter within the shell microstructure as the proportion of weight loss during the thermal treatment between 150°C and 550°C (fig. S2).

### Environmental characterization

We selected three key environmental drivers based on their known influence on mussel growth, their level of collinearity across the geographic scale investigated and the forecasted major ocean alterations under climate change (*16*). For each site, measurements of sea surface temperature, salinity and Chl-*a* concentration, the latter being used as a proxy for food supply (*24*), were generated using the Copernicus Marine Environment Monitoring Service (CMEMS) (http://marine.copernicus.eu/). These climate datasets are composed of high-resolution physical and biogeochemical assimilated (integration of observational and predicted information) daily data (*N* = 2,191 per parameter) (see data file S1). To provide a first order approximation of the water conditions prevailing during the near-maximum rates of shell deposition (*30*), we expressed parameters as mean May-October values averaged over the 6-year period 2009-2014 and used these as input variables (Fig. 1D, table S4).

Direct environmental monitoring for each site was not feasible due to the number of populations analysed, their geographic range (> 3,300 km) and the temporal resolution (6 years) required to estimate the average growth conditions during the lifespan of sampled specimens. For this large-scale study, remote-sensing and assimilated data presented potential advantages compared to traditional measurements due to their high spatio-temporal resolution, advanced calibration and validation (i.e. high correlation with discrete field measurements).

### Statistical analysis

GLMMs were applied to account for the hierarchical structure of the dataset consisting of multiple specimens (*N* = 24-26 replicates) from each collection site and to generalize our results to *Mytilus* populations beyond the study sample.

We carried out data exploration following the protocol of Zuur *et al.* (*39*). Initial inspection revealed no outliers. Pairwise scatterplots and variance inflation factors (VIFs) were calculated to check for collinearity between input variables. VIF values < 2 indicated an acceptable degree of correlation among covariates to be included within the same model. We applied residual regression to uncouple the unique from the shared contribution of temperature and Chl-*a* concentration to the response (*40*). This allowed us to account for the existing causal link between these two parameters and to avoid inferential problems from modelling non-independent covariates without losing explanatory power (*40*). To directly compare model estimates from predictors on different measurement scales, estimate biologically meaningful intercepts and interpret main effects when interactions are present, we standardised all the input variables (environmental parameters and shell length). For standardisation, we subtracted the sample mean from the variable values and divided them by the sample standard deviation 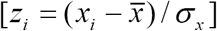. Preliminary inspection of residual patterns showed heteroscedasticity in most models. The use of different continuous probability distributions (i.e. gamma) and link functions did not stabilize the variance, therefore a ln-transformation of the response was required, except for calcite% and wt% measurements. Response variables did not require further transformations.

We used separate GLMMs to explore patterns of shell thickness of individual layers with latitude and shell length (size). The proportion (wt%) of organic matrix (*N* = 80) was modelled as a function of site (categorical, four levels) and prismatic thickness (continuous) to test for differences between polar and temperate regions and association with shell thickness. The response variable was coded as a value from 0 to 1; therefore, we used a GLM with a beta distribution and a logistic link function. Pair-wise contrasts with a Bonferroni correction were then used to test for differences in wt% among sites within and between climatic regions.

Different approaches were used to investigate the relationships between shell thickness and environmental gradients. Whole-shell thickness, periostracum thickness and calcite% were modelled separately (*N* = 424 each). Prismatic and nacreous layer thickness were analysed within the same GLMM, allowing the simultaneous prediction of common and divergent environmental effects on both layers and to reduce the probability of type I error. To model shell thickness (*N* = 424 × 2 layers) as a function of the environmental predictors we used a GLMM with a normal distribution (equation (1)). In the initial model, fixed continuous covariates were the standardised temperature, salinity and Chl-*a* in addition to shell layer (categorical, two levels) and their two-way interactions. Shell length (continuous) was included to control for possible effects of within population age on layer thickness. To incorporate the dependency among observations for a specific layer from the same collection site, we used site as a random intercept.

Models were optimized by first selecting the random structure and then the optimal fixed component. The principal tools for model comparison were the corrected Akaike Information Criterion (AICc) and bootstrapped likelihood ratio tests. Random terms were selected on prior knowledge of the dependency structure of the dataset. Visual inspection of residual patterns indicated violation of homogeneity in most cases. This required the use of variance structures (generalized least squares) allowing the residual spread to vary with respect to shell layer. The fixed component was optimized by rejecting only non-significant interaction terms that minimized the AICc value. For all model comparisons, variation of AICc between the optimal (lowest AICc value) and competing models were greater than 8, and fixed-effect estimates were nearly identical, indicating that competing models were very unlikely to be superior (*41*). The proportion of variance explained by the models was quantified with conditional or pseudo determination coefficients (cR^2^ or pseudoR^2^). We used variograms to assess the absence of spatial autocorrelation. Final models were validated by inspection of standardised residual patterns to verify GLMM assumptions of normality, homogeneity and independence. We used optimal models to estimate the mean effect sizes (same measurement scale) of environmental drivers on the response. Ninety-five per cent confidence intervals (95% CI) for the regression parameters were generated using bias-corrected parametric bootstrap methods (10,000 iterations). 95% CIs were used for statistical inference due to estimation of approximated significance values (*P*-value) in mixed-modelling. If the confidence intervals did not overlap zero, then the effect was considered significant. All data exploration and statistical analyses were performed in R (v3.4.1) (for packages see table S5).

## Acknowledgments

We thank Iain Johnston (Scottish Oceans Institute, St. Andrews, UK), Sarah Dashfield (Plymouth Marine Laboratory, Plymouth, UK), Dr. Peter Thor (Norwegian Polar Institute, Tromsø, Norway), Alexander Ventura (University of Gothenburg, Kristineberg, Sweden), Dr. Henk van der Veer and Rob Dekker (Royal Netherlands Institute for Sea Research, Texel, Netherlands) for help with specimens collection, and the Statistics Clinic (University of Cambridge, UK) for statistical advice.

## Author contributions

L.T., L.S.P. and E.M.H. conceived the original project and designed the study; L.T. performed laboratory work and thermogravimetric analysis, generated environmental datasets, performed modelling work and analysed output data; L.T., T.S., J.T. and M.K.S. collected materials; L.T., L.S.P. and E.M.H. wrote the first draft of the manuscript, and all co-authors contributed substantially to revisions.

## SUPPLEMENTARY MATERIALS

**Fig. S1.**
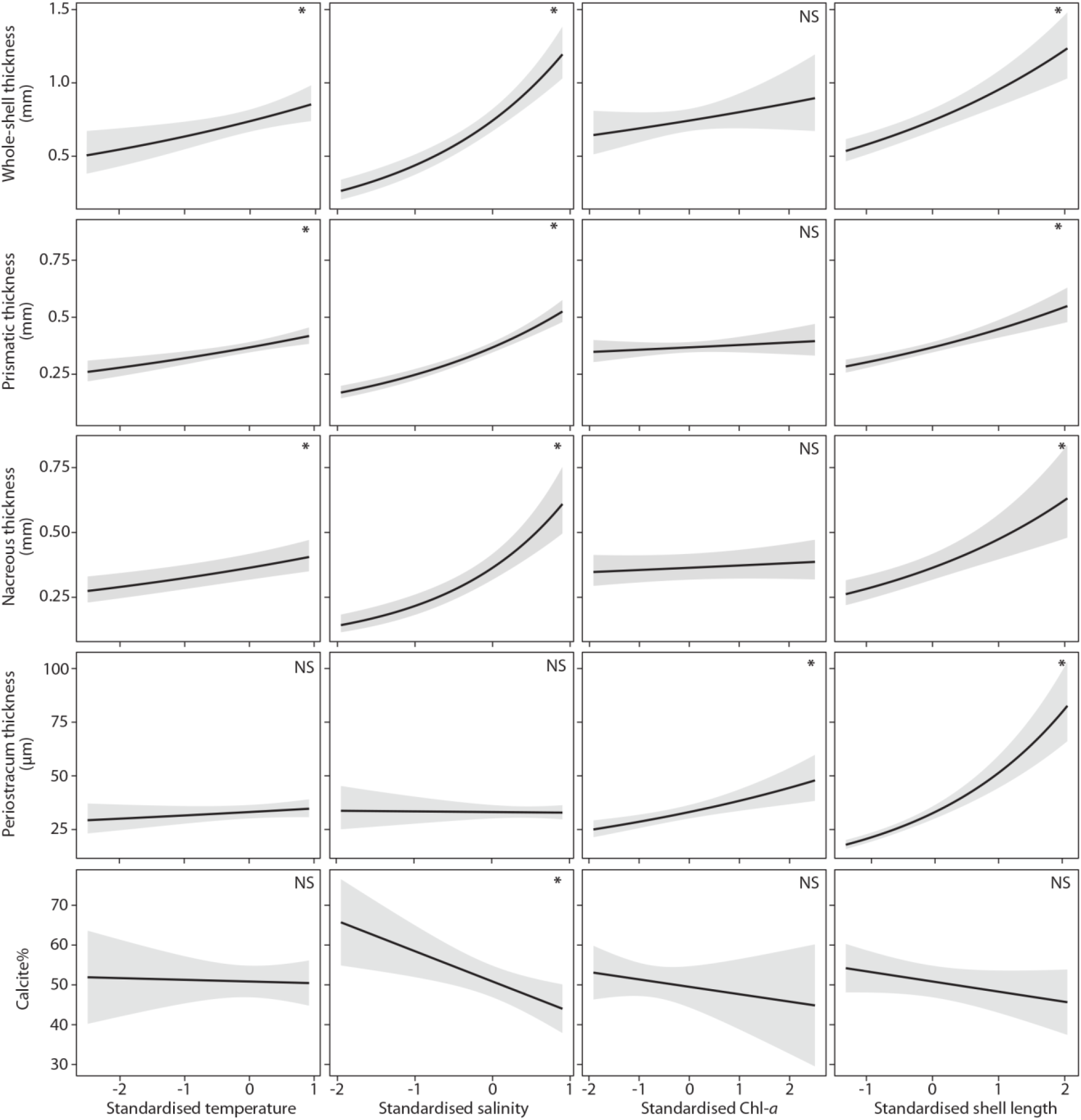
Relationships between shell layers and modelled predictors. Predicted relationships between the whole-shell, prismatic layer, nacreous layer, periostracum thickness and the calcite% with standardised water temperature [mean (SD) = 13.03°C (4.32)], salinity [mean (SD) = 25.52 psu (10.29)], Chl-*a* concentration [mean (SD) = 2.48 mg m^-3^ (1.41)], and shell length [mean (SD) = 47.42 mm (16.20)]. Predicted values (continuous lines) and confidence intervals (shaded areas) across the range of each predictor were estimated controlling statistically for the other covariates (mean values). (NS *P* > 0.05, * *P* < 0.05)

**Fig. S2.**
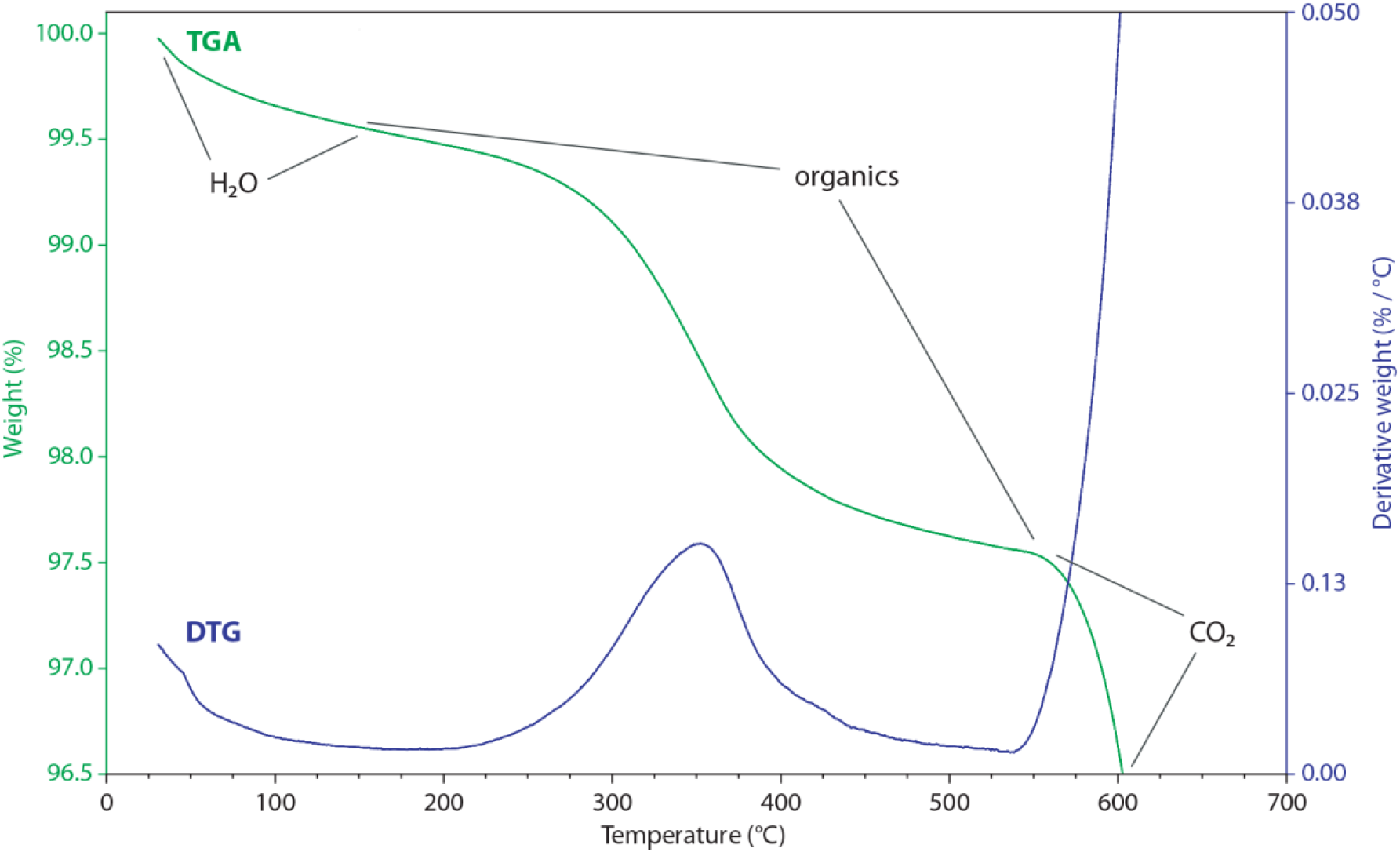
Example of Thermogravimetric Analysis (TGA) and Derivative Thermogravimetry (DTG). The TGA curve (green) represents the weight changes with increasing treatment temperature for the prismatic layer of *Mytilus edulis*. The sample was exposed to a constant heating, from ∼25°C to 700°C at a linear rate of 10°C min^-1^. Three known regions of weight loss with increasing temperature are highlighted (*42*): i) the evaporation of physically adsorbed water at 30-150°C, ii) the degradation of organics at 150-550 °C, and iii) the rapid decomposition of calcium carbonate (CaCO_3_) into calcium oxide (CaO) and carbon dioxide (CO_2_) starting at ∼550°C. The DTG line (blue) represents the derivative of the thermal curve and shows the rate of weight loss during heating. The peak indicates the temperature at which the organic mass loss was fastest.

**Table S1.**
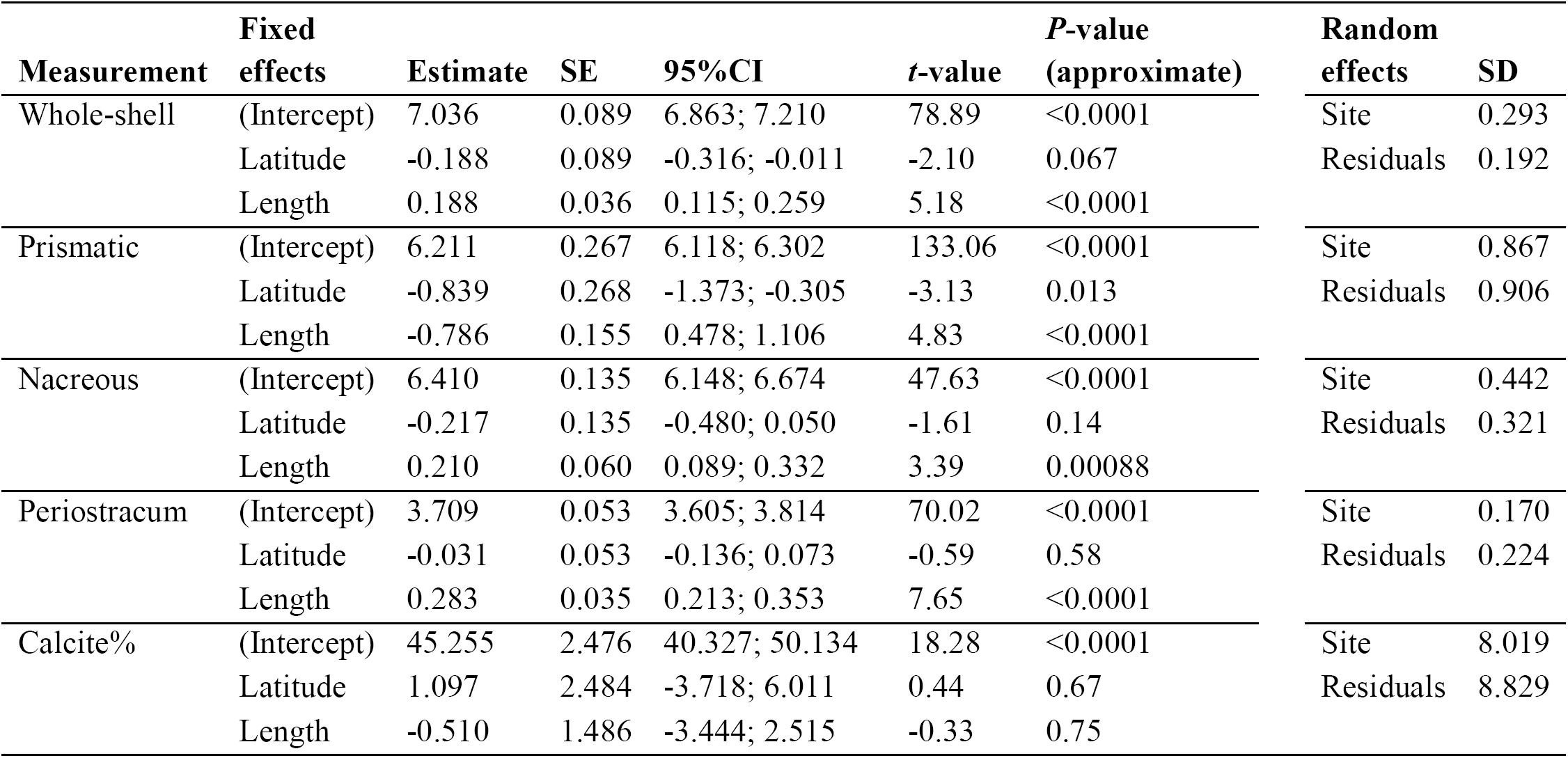
Latitudinal GLMMs summary. Estimated statistics and bootstrapped 95% CI for regression parameters are reported for the modelled relationships between individual shell measurements, standardised latitude and shell length.

| Measurement | Fixed effects | Estimate | SE | 95%CI | t-value | P-value (approximate) | Random effects | SD |
| --- | --- | --- | --- | --- | --- | --- | --- | --- |
| Whole-shell | (Intercept) | 7.036 | 0.089 | 6.863; 7.210 | 78.89 | <0.0001 | Site | 0.293 |
|  | Latitude | -0.188 | 0.089 | -0.316; -0.011 | -2.10 | 0.067 | Residuals | 0.192 |
|  | Length | 0.188 | 0.036 | 0.115; 0.259 | 5.18 | <0.0001 |  |  |
| Prismatic | (Intercept) | 6.211 | 0.267 | 6.118; 6.302 | 133.06 | <0.0001 | Site | 0.867 |
|  | Latitude | -0.839 | 0.268 | -1.373; -0.305 | -3.13 | 0.013 | Residuals | 0.906 |
|  | Length | -0.786 | 0.155 | 0.478; 1.106 | 4.83 | <0.0001 |  |  |
| Nacreous | (Intercept) | 6.410 | 0.135 | 6.148; 6.674 | 47.63 | <0.0001 | Site | 0.442 |
|  | Latitude | -0.217 | 0.135 | -0.480; 0.050 | -1.61 | 0.14 | Residuals | 0.321 |
|  | Length | 0.210 | 0.060 | 0.089; 0.332 | 3.39 | 0.00088 |  |  |
| Periostracum | (Intercept) | 3.709 | 0.053 | 3.605; 3.814 | 70.02 | <0.0001 | Site | 0.170 |
|  | Latitude | -0.031 | 0.053 | -0.136; 0.073 | -0.59 | 0.58 | Residuals | 0.224 |
|  | Length | 0.283 | 0.035 | 0.213; 0.353 | 7.65 | <0.0001 |  |  |
| Calcite% | (Intercept) | 45.255 | 2.476 | 40.327; 50.134 | 18.28 | <0.0001 | Site | 8.019 |
|  | Latitude | 1.097 | 2.484 | -3.718; 6.011 | 0.44 | 0.67 | Residuals | 8.829 |
|  | Length | -0.510 | 1.486 | -3.444; 2.515 | -0.33 | 0.75 |  |  |

**Table S2.** Environmental GLMMs summary. Estimated statistics and bootstrapped 95% CI for regression parameters are reported for the modelled relationships between individual shell measurements, standardised environmental covariates and shell length.

| Measurement | Fixed effects | Estimate | SE | 95%CI | t-value | P-value (approximate) | Random effects | SD |
| --- | --- | --- | --- | --- | --- | --- | --- | --- |
| Whole-shell | (Intercept) | 6.617 | 0.051 | 6.517; 6.717 | 128.62 | <0.0001 | Site | 0.209 |
|  | Temperature | 0.156 | 0.054 | 0.014; 0.240 | 2.89 | 0.013 | Residual | 0.188 |
|  | Salinity | 0.525 | 0.060 | 0.411; 0.672 | 8.69 | <0.0001 |  |  |
|  | Chl-*a* | 0.074 | 0.054 | -0.042; 0.216 | 1.37 | 0.20 |  |  |
|  | Length | 0.248 | 0.037 | 0.181; 0.327 | 6.44 | <0.0001 |  |  |
| Prismatic (Pr)<br>and nacreous<br>(Na) | (Intercept) | 5.907 | 0.031 | 5.774; 6.036 | 188.31 | <0.0001 | Site (Pr) | 0.123 |
|  | Temperature | 0.138 | 0.033 | 0.013; 0.260 | 4.17 | 0.0011 | Site (Na) | 0.310 |
|  | Salinity | 0.396 | 0.039 | 0.264; 0.531 | 10.22 | <0.0001 | Residual | 0.198 |
|  | Chl-*a* | 0.028 | 0.033 | -0.087; 0.139 | 0.86 | 0.41 |  |  |
|  | Length | 0.197 | 0.031 | 0.096; 0.297 | 6.39 | <0.0001 |  |  |
|  | Layer(Na) | -0.054 | 0.078 | -0.247; 0.134 | -0.70 | 0.49 |  |  |
|  | Salinity ×<br>Layer(Na) | 0.259 | 0.088 | 0.065; 0.443 | 2.95 | 0.0033 |  |  |
|  | Length ×<br>Layer(Na) | 0.111 | 0.067 | -0.036; 0.258 | 1.66 | 0.096 |  |  |
| Periostracum | (Intercept) | 3.500 | 0.048 | 3.406; 3.596 | 71.03 | <0.0001 | Site | 0.130 |
|  | Temperature | 0.049 | 0.043 | -0.036; 0.134 | 1.12 | 0.28 | Residual | 0.230 |
|  | Salinity | -0.009 | 0.061 | -0.131; 0.111 | -0.14 | 0.89 |  |  |
|  | Chl-*a* | 0.147 | 0.038 | 0.071; 0.221 | 3.88 | 0.002 |  |  |
|  | Length | 0.439 | 0.041 | 0.357; 0.522 | 10.25 | <0.0001 |  |  |
|  | Temperature ×<br>Length | -0.064 | 0.035 | -0.135; 0.006 | -1.77 | 0.082 |  |  |
|  | Salintiy ×<br>Length | -0.151 | 0.061 | -0.271; -0.029 | -2.38 | 0.020 |  |  |
| Calcite% | (Intercept) | 51.092 | 1.903 | 47.338; 54.927 | 26.84 | <0.0001 | Site | 7.605 |
|  | Temperature | -4.003 | 2.112 | -4.508; 3.791 | -0.19 | 0.85 | Residual | 9.656 |
|  | Salinity | -7.163 | 2.476 | -12.025; -2.382 | -2.87 | 0.012 |  |  |
|  | Chl-*a* | -1.908 | 2.392 | -6.662; 2.764 | -0.80 | 0.44 |  |  |
|  | Length | -2.404 | 1.690 | -5.807; 1.071 | -1.367 | 0.18 |  |  |

**Table S3.**
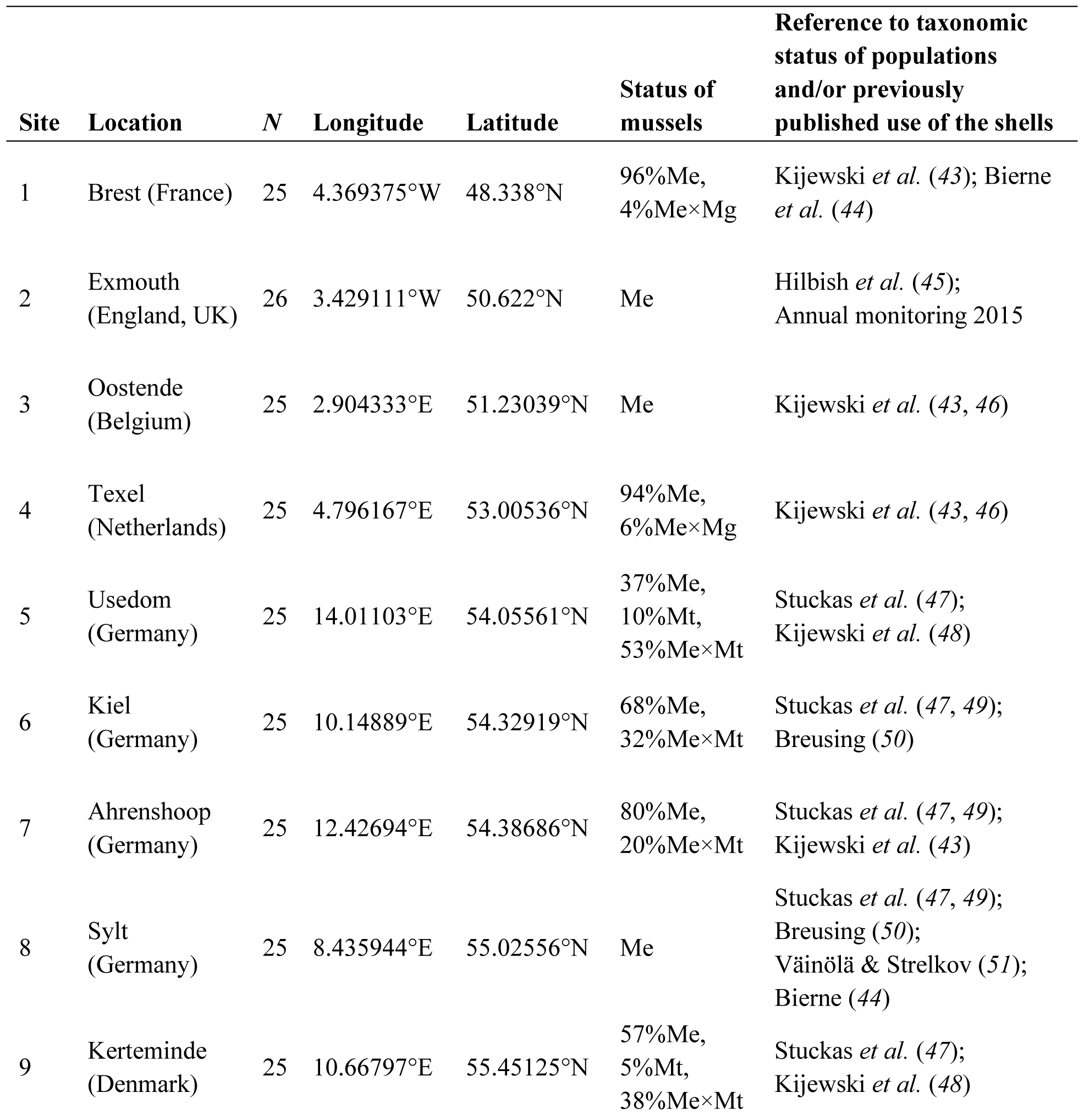
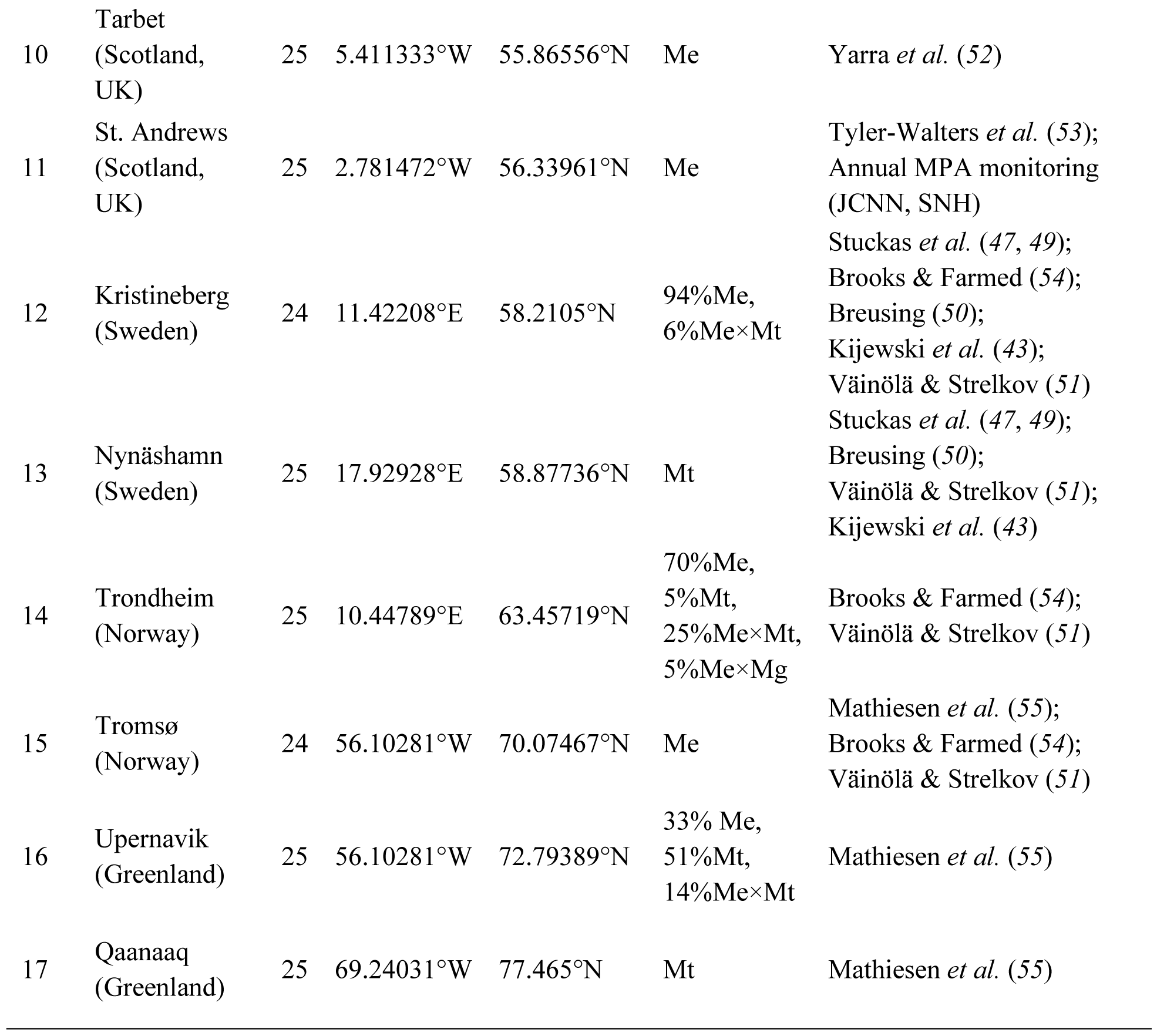
Provenience and taxonomic status of the *Mytilus* populations used for the study. For each sampling site (site codes as in Fig. 1C), geographic location, samples size (*N*), site coordinates (longitude and latitude), genotypic status [proportion of *Mytilus edulis* (Me), *M. trossulus* (Mt), *M. galloprovincialis* (Mg) and hybrids] and reference and/or previous use of the studied populations are reported.

| Site | Location | *N* | Longitude | Latitude | Status of mussels | Reference to taxonomic status of populations and/or previously published use of the shells |
| --- | --- | --- | --- | --- | --- | --- |
| 1 | Brest (France) | 25 | 4.369375°W | 48.338°N | 96% Me,<br>4% Me×Mg | Kijewski *et al.* (43); Bierne *et al.* (44) |
| 2 | Exmouth (England, UK) | 26 | 3.429111°W | 50.622°N | Me | Hilbish *et al.* (45);<br>Annual monitoring 2015 |
| 3 | Oostende (Belgium) | 25 | 2.904333°E | 51.23039°N | Me | Kijewski *et al.* (43, 46) |
| 4 | Texel (Netherlands) | 25 | 4.796167°E | 53.00536°N | 94% Me,<br>6% Me×Mg | Kijewski *et al.* (43, 46) |
| 5 | Usedom (Germany) | 25 | 14.01103°E | 54.05561°N | 37% Me,<br>10% Mt,<br>53% Me×Mt | Stuckas *et al.* (47);<br>Kijewski *et al.* (48) |
| 6 | Kiel (Germany) | 25 | 10.14889°E | 54.32919°N | 68% Me,<br>32% Me×Mt | Stuckas *et al.* (47, 49);<br>Breusing (50) |
| 7 | Ahrenschoop (Germany) | 25 | 12.42694°E | 54.38686°N | 80% Me,<br>20% Me×Mt | Stuckas *et al.* (47, 49);<br>Kijewski *et al.* (43) |
| 8 | Sylt (Germany) | 25 | 8.435944°E | 55.02556°N | Me | Stuckas *et al.* (47, 49);<br>Breusing (50);<br>Väinölä & Strelkov (51);<br>Bierne (44) |
| 9 | Kerteminde (Denmark) | 25 | 10.66797°E | 55.45125°N | 57% Me,<br>5% Mt,<br>38% Me×Mt | Stuckas *et al.* (47);<br>Kijewski *et al.* (48) |
| 10 | Tarbet<br>(Scotland,<br>UK) | 25 | 5.411333°W | 55.86556°N | Me | Yarra *et al.* (52) |
| 11 | St. Andrews<br>(Scotland,<br>UK) | 25 | 2.781472°W | 56.33961°N | Me | Tyler-Walters *et al.* (53);<br>Annual MPA monitoring<br>(JCNN, SNH) |
| 12 | Kristineberg<br>(Sweden) | 24 | 11.42208°E | 58.2105°N | 94% Me,<br>6% Me×Mt | Stuckas *et al.* (47, 49);<br>Brooks & Farmed (54);<br>Breusing (50);<br>Kijewski *et al.* (43);<br>Väinölä & Strelkov (51) |
| 13 | Nynäshamn<br>(Sweden) | 25 | 17.92928°E | 58.87736°N | Mt | Stuckas *et al.* (47, 49);<br>Breusing (50);<br>Väinölä & Strelkov (51);<br>Kijewski *et al.* (43) |
| 14 | Trondheim<br>(Norway) | 25 | 10.44789°E | 63.45719°N | 70% Me,<br>5% Mt,<br>25% Me×Mt,<br>5% Me×Mg | Brooks & Farmed (54);<br>Väinölä & Strelkov (51) |
| 15 | Tromsø<br>(Norway) | 24 | 56.10281°W | 70.07467°N | Me | Mathiesen *et al.* (55);<br>Brooks & Farmed (54);<br>Väinölä & Strelkov (51) |
| 16 | Upernavik<br>(Greenland) | 25 | 56.10281°W | 72.79389°N | 33% Me,<br>51% Mt,<br>14% Me×Mt | Mathiesen *et al.* (55) |
| 17 | Qaanaaq<br>(Greenland) | 25 | 69.24031°W | 77.465°N | Mt | Mathiesen *et al.* (55) |

**Table S4.**
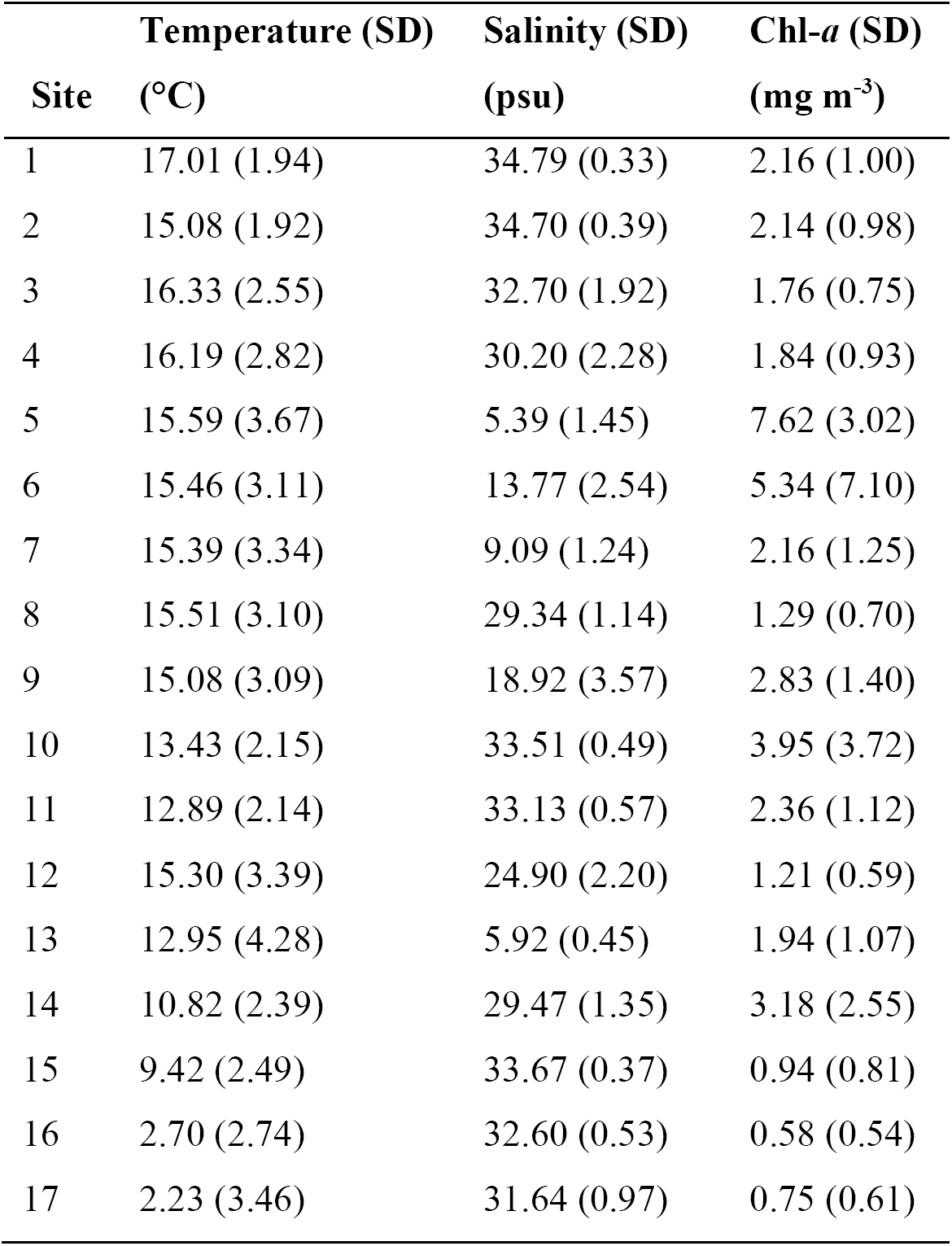
Environmental covariates. Summary statistics (mean value and SD) of environmental conditions at each study site. Site codes as in Fig. 1C.

|  | Temperature (SD) | Salinity (SD) | Chl-*a* (SD) |
| --- | --- | --- | --- |
| Site | (°C) | (psu) | (mg m<sup>-3</sup>) |
| 1 | 17.01 (1.94) | 34.79 (0.33) | 2.16 (1.00) |
| 2 | 15.08 (1.92) | 34.70 (0.39) | 2.14 (0.98) |
| 3 | 16.33 (2.55) | 32.70 (1.92) | 1.76 (0.75) |
| 4 | 16.19 (2.82) | 30.20 (2.28) | 1.84 (0.93) |
| 5 | 15.59 (3.67) | 5.39 (1.45) | 7.62 (3.02) |
| 6 | 15.46 (3.11) | 13.77 (2.54) | 5.34 (7.10) |
| 7 | 15.39 (3.34) | 9.09 (1.24) | 2.16 (1.25) |
| 8 | 15.51 (3.10) | 29.34 (1.14) | 1.29 (0.70) |
| 9 | 15.08 (3.09) | 18.92 (3.57) | 2.83 (1.40) |
| 10 | 13.43 (2.15) | 33.51 (0.49) | 3.95 (3.72) |
| 11 | 12.89 (2.14) | 33.13 (0.57) | 2.36 (1.12) |
| 12 | 15.30 (3.39) | 24.90 (2.20) | 1.21 (0.59) |
| 13 | 12.95 (4.28) | 5.92 (0.45) | 1.94 (1.07) |
| 14 | 10.82 (2.39) | 29.47 (1.35) | 3.18 (2.55) |
| 15 | 9.42 (2.49) | 33.67 (0.37) | 0.94 (0.81) |
| 16 | 2.70 (2.74) | 32.60 (0.53) | 0.58 (0.54) |
| 17 | 2.23 (3.46) | 31.64 (0.97) | 0.75 (0.61) |

**Table S5.**
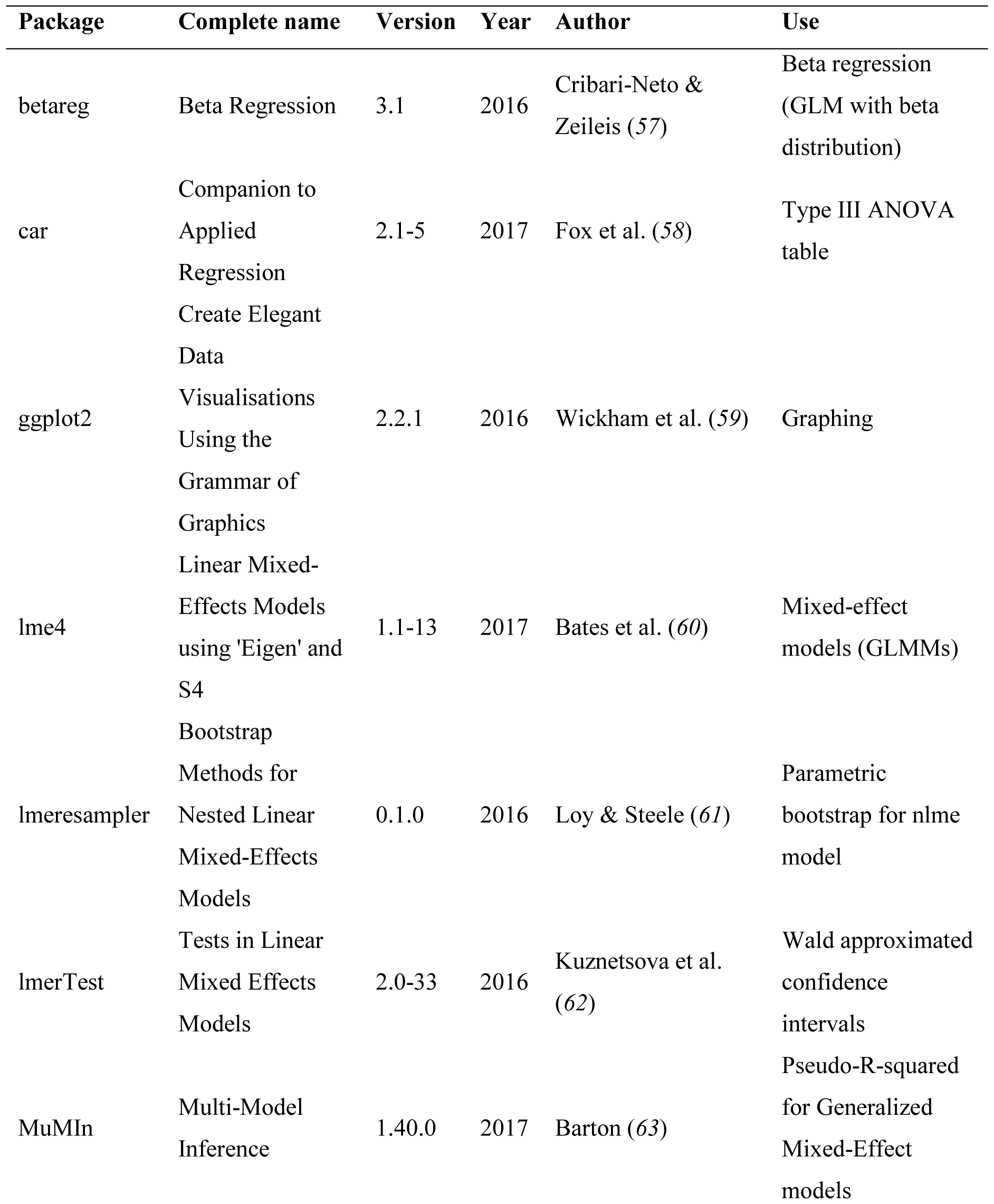
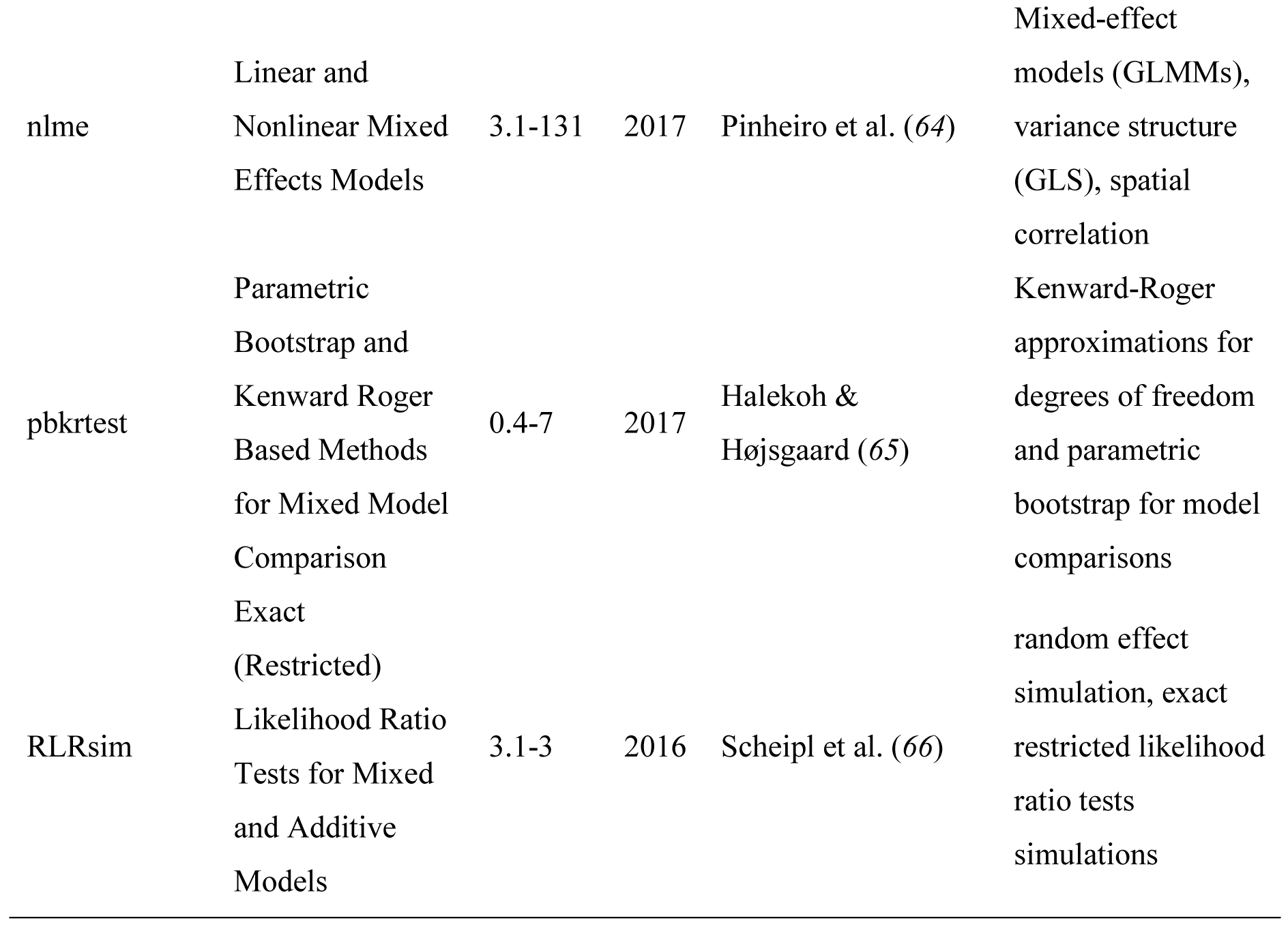
List of R packages. Packages and version used with the R software (v3.4.1) (*56*) for data exploration, statistical analysis and graphing.

## SUPPLEMENTARY MATERIALS AND METHODS

### Protocol for Thermal Gravimetric Analyses

Thermogravimetric Analyses (TGA) are reported following the guidelines made by the Committee on Standardisation of the International Confederation for Thermal Analysis and Calorimetry (ICTAC) and appeared in standards as ASTM E 472 (1991) (*67*, *68*).

A. **Properties of the sample**
  1. *Source of material and identification*
    - Shell of wild Atlantic blue mussel (*Mytilus edulis*).
    - Prismatic layer composed of calcium carbonate (CaCO_3_, calcite), variable amount of organics (∼1-2%) and trace elements, such as quartzite (SiO_2_) and magnesium (Mg).
  2. *Sample history*
    - Shells were cleaned, raised with mill-Q water, dried at room temperature for seven days.
    - The periostracum was removed by sanding and a tile of prismatic layer isolated (8 × 5 mm) with a Dremel rotary tool (Dremel 300/395RD MultiPro, Racine, Wisconsin, USA).
    - Samples were cleaned in an ultrasonic bath (Ultrasonic Cleaner CD-4800, Practical Systems Inc., Odessa, FL, USA) with mill-Q water, air-dried and powdered with an agate mortar.
    - Additional, oven drying (30°C for 24h, convection oven) to remove residual pre-treatment water.
  3. *Physical properties*
    - Fine grade powder.

A. **Experimental conditions**
  1. *Apparatus used*
    - Thermogravimetric Analyser: TGA Q500, TA instrument (New Castle, DE, USA) Q series.
  2. *Thermal treatment*
    - Initial temperature, ∼25°C (room temperature).
    - Final temperature, 700°C.
    - Linear rate of heating, 10°C min^-1^.
  3. *Sample atmosphere*
    - Dynamic (flowing) atmosphere.
    - Flow rate for balance 40 ml min^-1^ and for sample 60ml min^-1^.
    - Gas composition: nitrogen, “white spot”.
  4. *Sample holder*
    - Platinum crucible, cylindrical: diameter 10 mm and height 1.5 mm.
    - Sample was tipped and spread to cover the bottom of the crucible.
  5. *Sample mass*
    - 10 mg of powder were weighted on a separate micro-balance (Ultramicro 4504 MP8, Sartorius, Göttingen; readability 0.1 µg).

A. **Data acquisition and manipulation methods**
  1. *Software version*
    - Universal Analysis 2000, version 4.5A, TA instrument (New Castle, DE, USA).

## SUPPLEMENTARY DATA

### Environmental datasets

List of the datasets used for the calculation of mean annual values of environmental descriptors. Water temperature, salinity and chlorophyll-a concentrations were expressed as mean values (May - October) averaged over the 6-year period 2009 - 2014. This study has been conducted using the Copernicus Marine Service Products: COPERNICUS - Marine Environment Monitoring System (http://marine.copernicus.eu/).

## DATASET #1

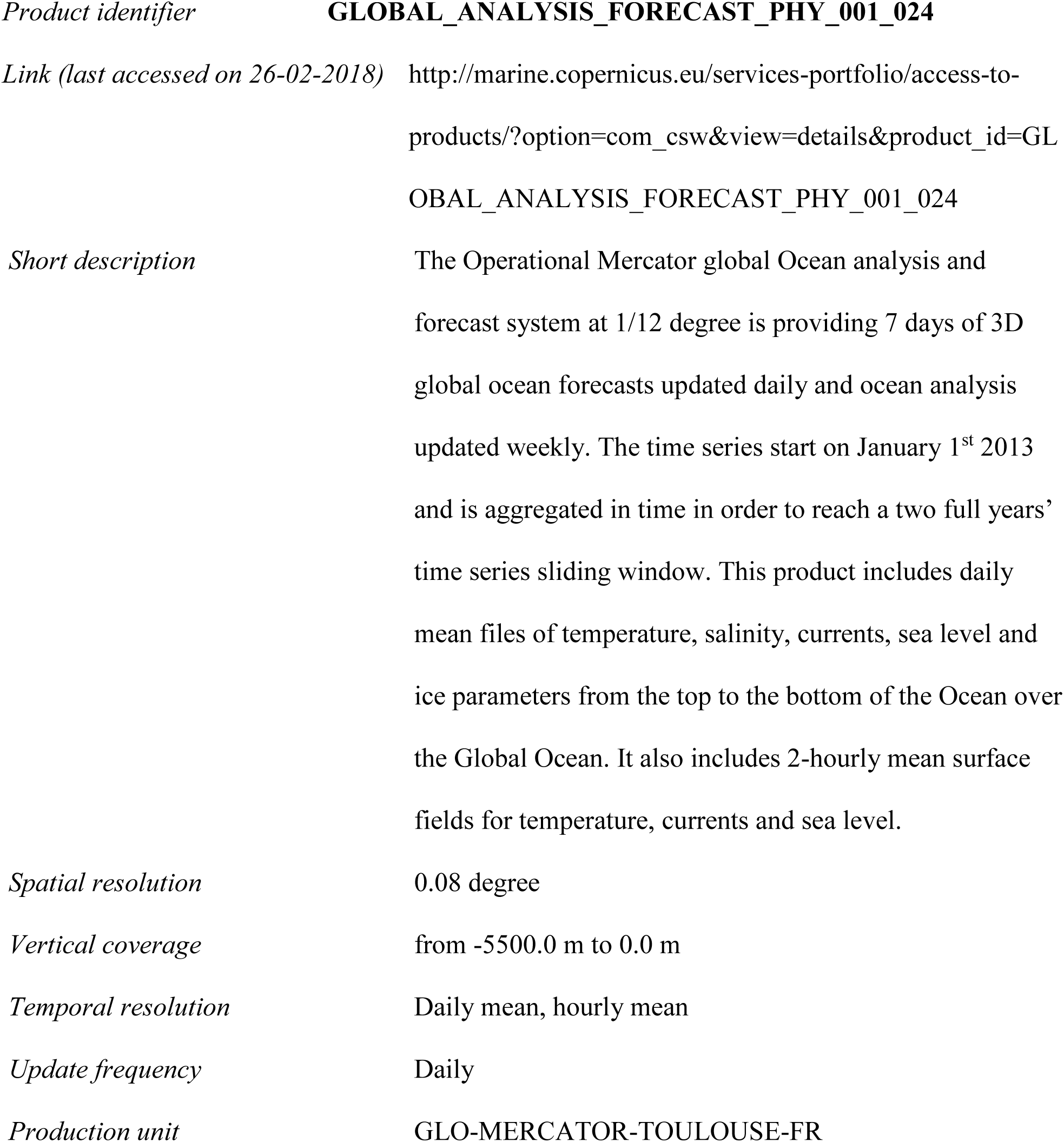

## DATASET #2

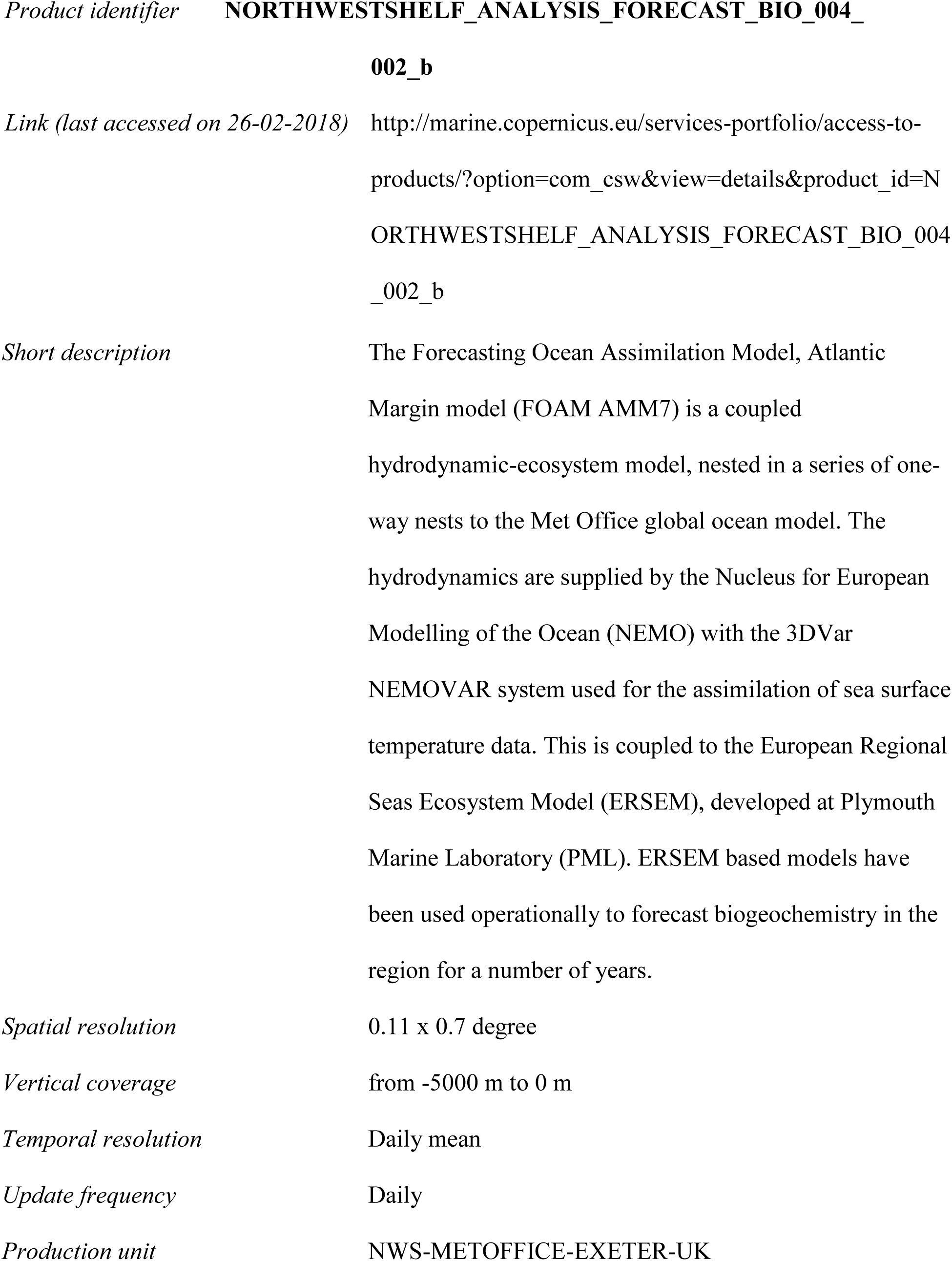

## DATASET #3

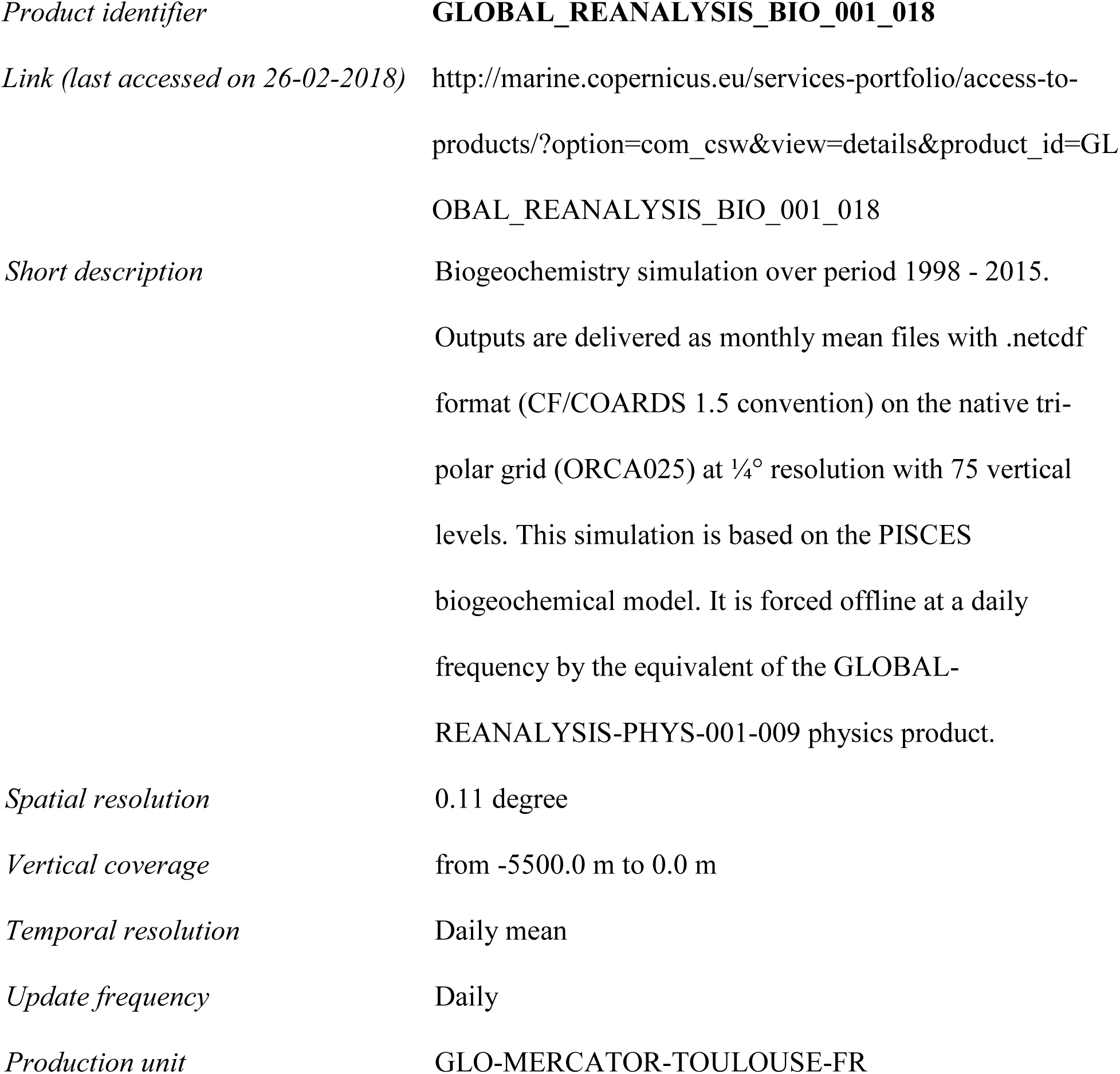

## DATASET #3

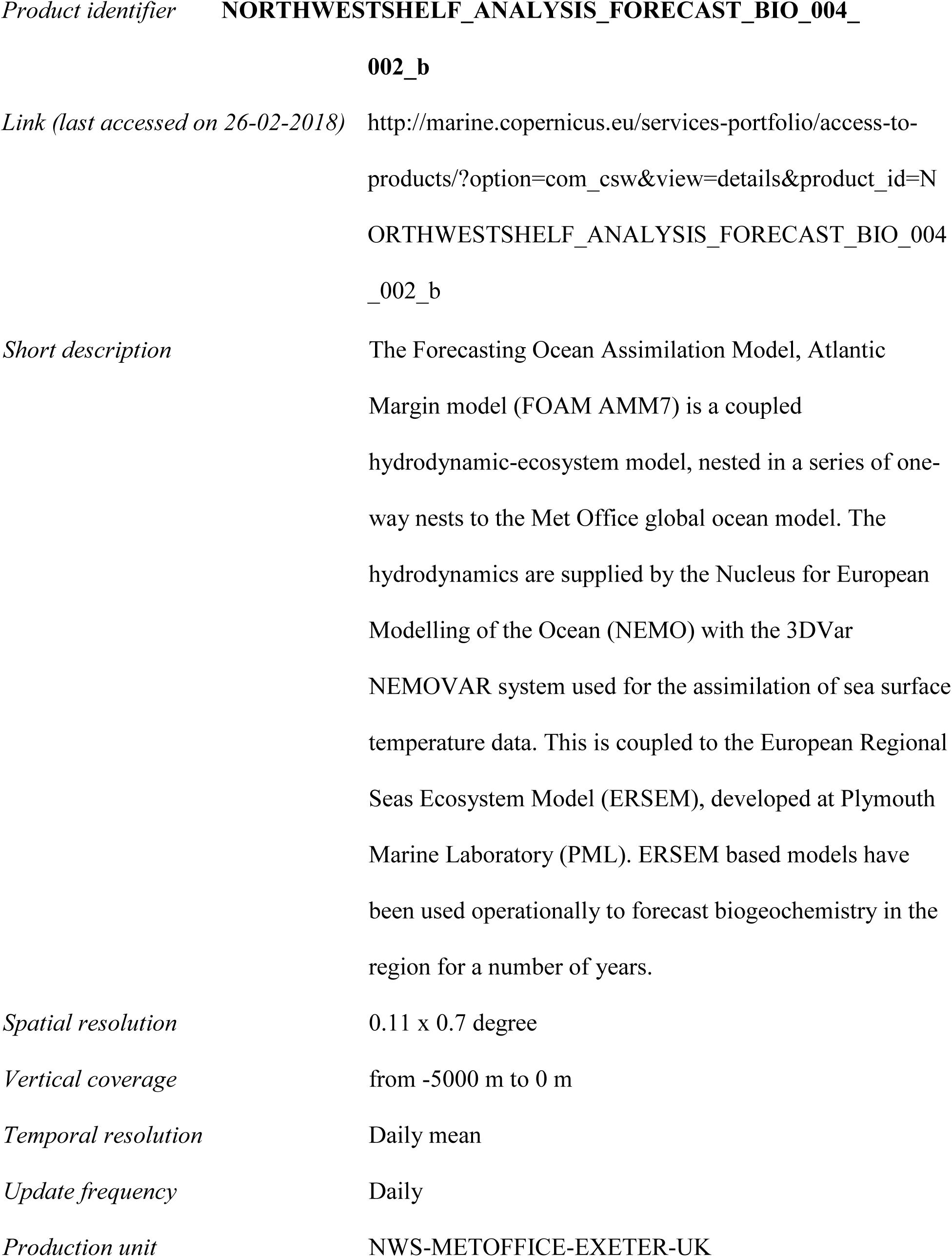

## DATASET #4

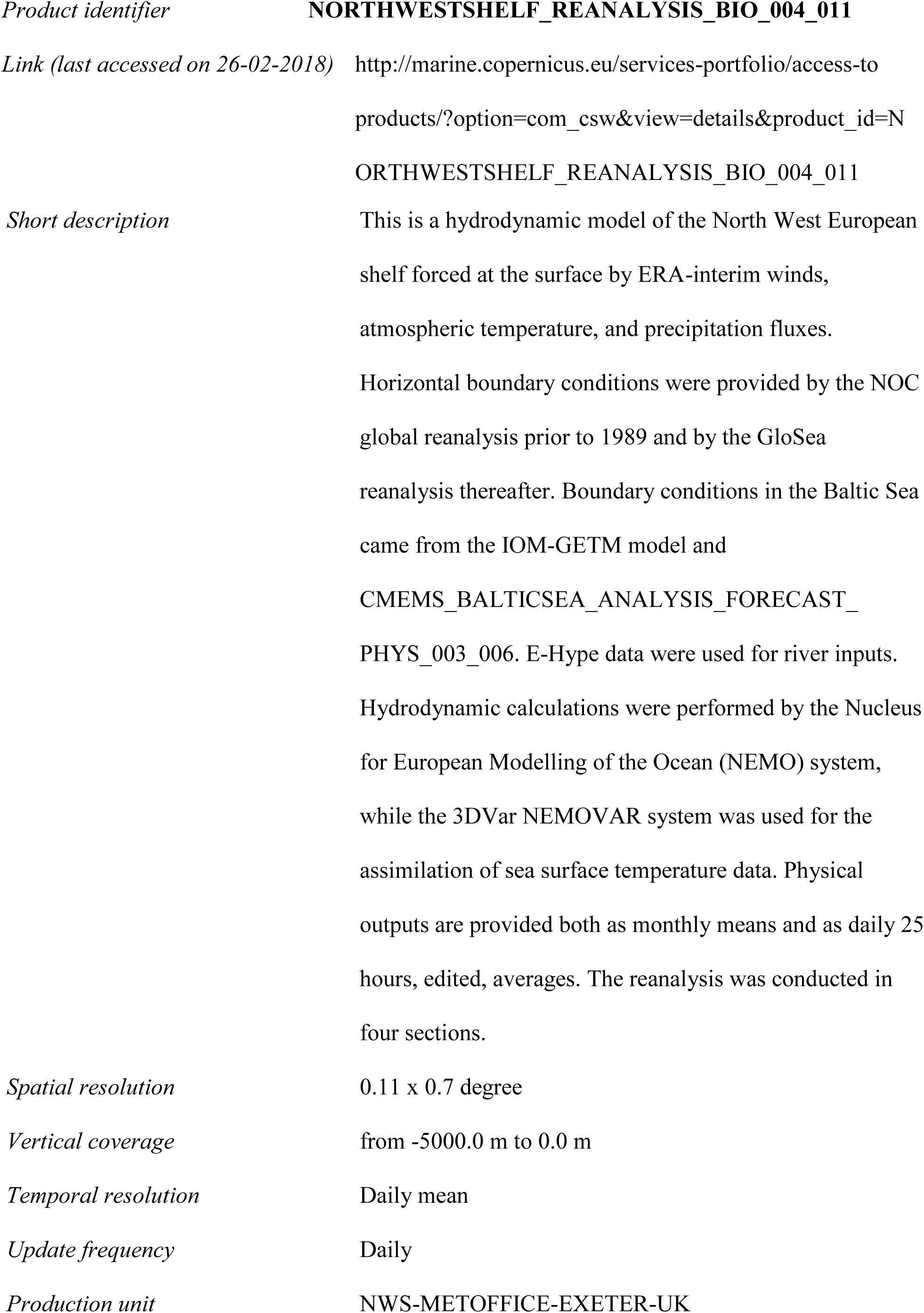

## DATASET #5

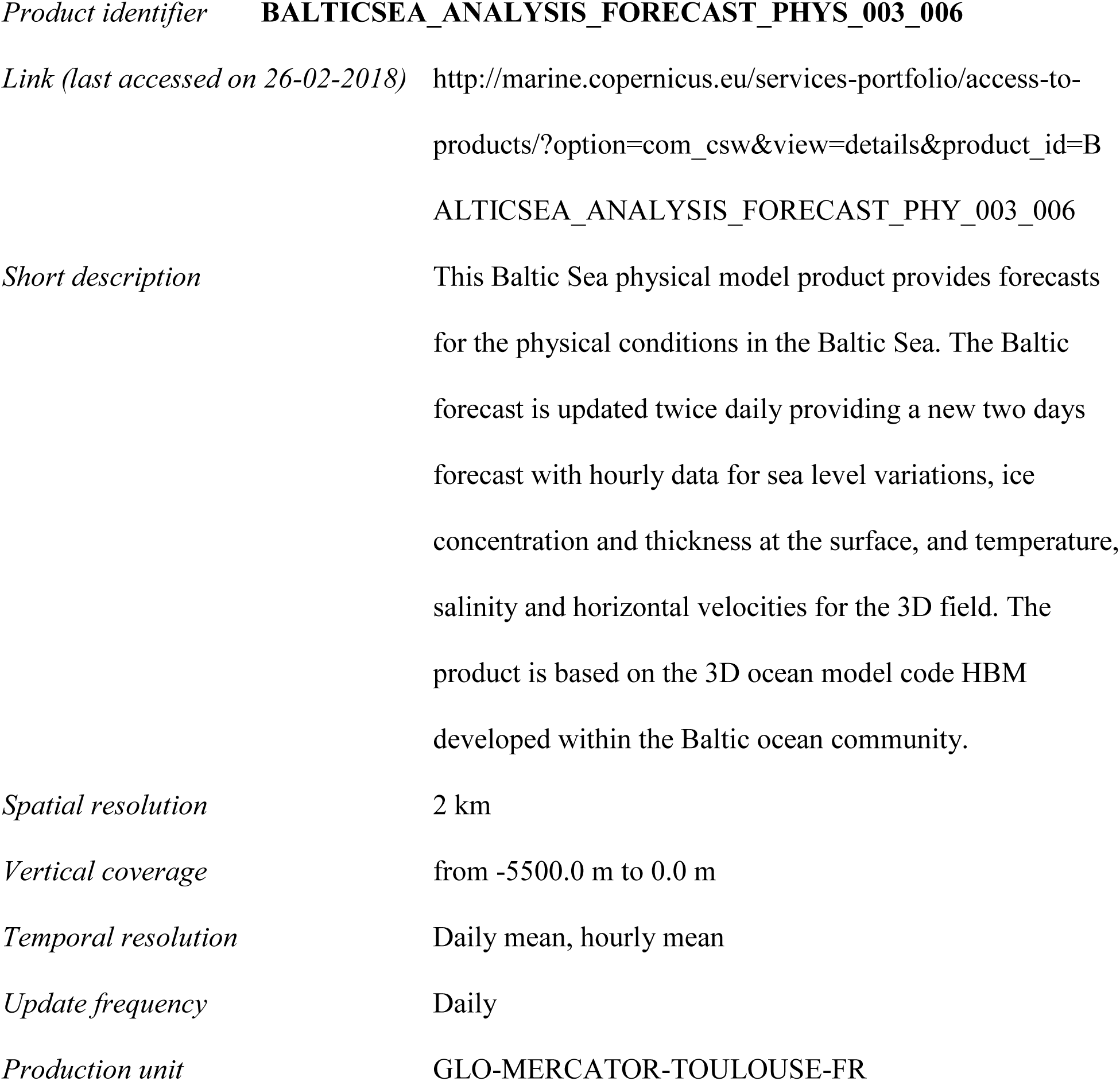

## DATASET #6

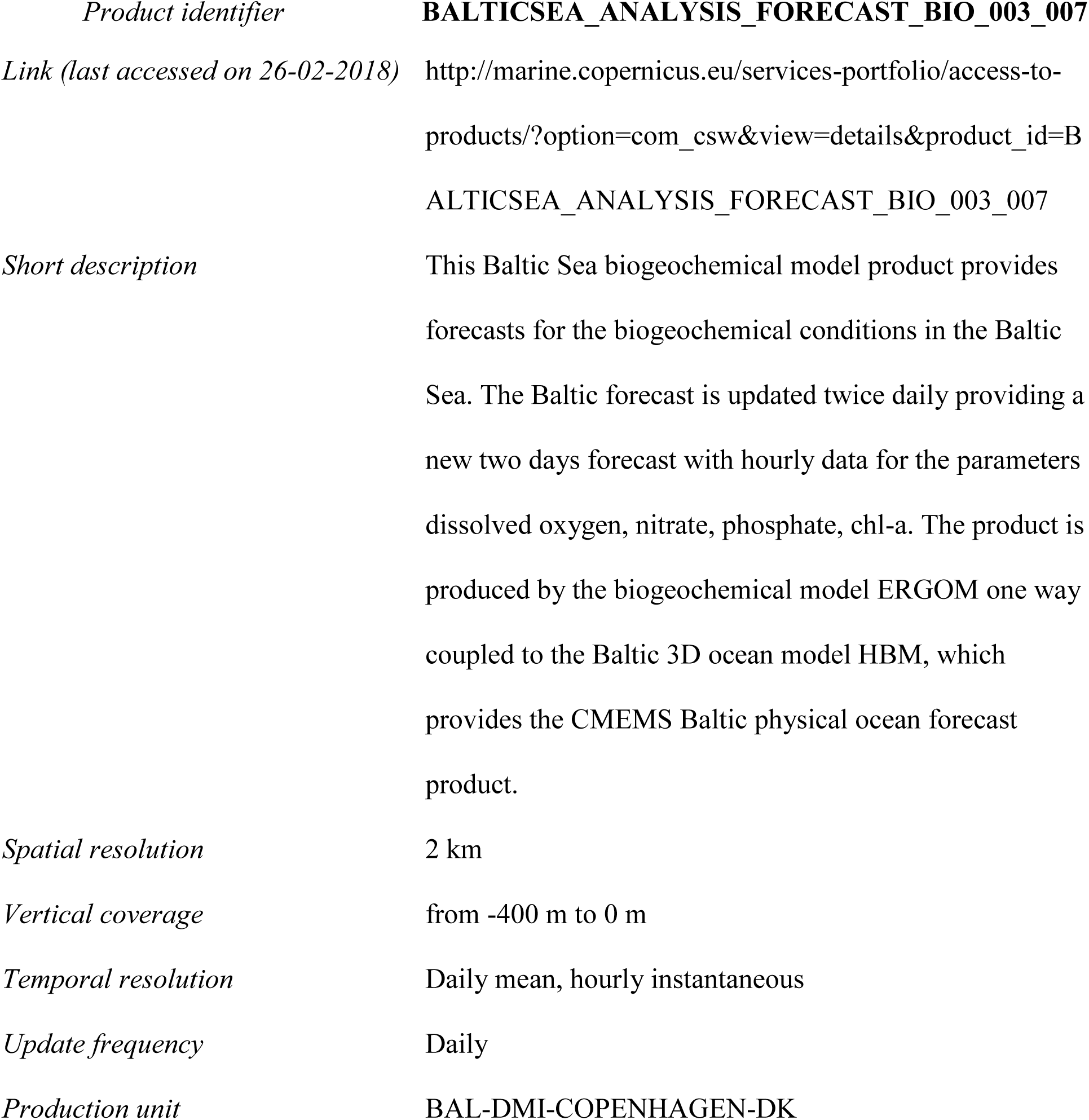

